# Persistent genomic erosion in whooping cranes despite demographic recovery

**DOI:** 10.1101/2024.12.12.628160

**Authors:** Claudia Fontsere, Samuel A. Speak, Andrew J. Caven, Juan Antonio Rodríguez, Xuejing Wang, Carolina Pacheco, Molly Cassatt-Johnstone, Georgette Femerling, Brigid Maloney, Jennifer Balacco, Joanna Collins, Ying Sims, Linelle Abueg, Olivier Fedrigo, Erich D. Jarvis, Barry K. Hartup, Beth Shapiro, M. Thomas P. Gilbert, Cock van Oosterhout, Hernán E. Morales

**Affiliations:** Globe Institute, University of Copenhagen, Denmark; School of Environmental Sciences, University of East Anglia, UK; Natural History Museum, London, UK; North of England Zoological Society, Chester Zoo, Chester, UK; International Crane Foundation, Baraboo, Wisconsin, USA; Crane Trust, Wood River, Nebraska, USA; Biology department, Lund University, Sweden; Department of Ecology and Evolutionary Biology, University of California Santa Cruz, USA; Department of Human Genetics, McGill University, Montreal, Quebec, Canada; Vertebrate Genome Lab, The Rockefeller University, New York, USA; School of Veterinary Medicine, University of Wisconsin-Madison, Wisconsin, USA; University Museum, NTNU, Trondheim, Norway

## Abstract

Integrating *in-situ* (wild) and *ex-situ* (captive) conservation efforts can mitigate genetic diversity loss and help prevent extinction of endangered wild populations. The whooping crane (*Grus americana*) experienced severe population declines in the 18th century, culminating into a collapse to 16 individuals in 1941. Legal protections and conservation actions have since increased the population to approximately 840 individuals, yet the impact on genomic diversity remains unclear. We analysed the temporal dynamics of genomic erosion by sequencing a high-quality genome reference, and re-sequencing 16 historical and 37 modern genomes, including wild individuals and four generations of captive-bred individuals. Genomic demographic reconstructions reveal a steady decline, accelerating over the past 300 years with the European settlement of North America. Temporal genomic analyses show that despite demographic recovery, the species has lost 70% of its genetic diversity and has increased their inbreeding. Although the modern population bottleneck reduced the ancestral genetic load, modern populations possess more realized load than masked load, possibly resulting in a chronic loss of fitness. Integrating pedigree and genomic data, we underscore the role of breeding management in reducing recent inbreeding. Yet ongoing heterozygosity loss, load accumulation, and background inbreeding argues against the species’ downlisting from their current Endangered status on the IUCN Red List and the Endangered Species Act. The presence of private genetic variation in wild and captive populations suggests that wild-captive crosses could enhance genetic diversity and reduce the realized load. Our findings emphasize the role of genomics in informing conservation management and policy.

## Introduction

The ongoing sixth mass extinction is reducing population sizes and driving species extinctions (Barnosky et al., 2011). Conservation strategies, including *in-situ* (wild) and *ex-situ* (captive) approaches, aim to restore populations and maintain genetic diversity (Braverman, 2014). The One Plan Approach integrates these strategies, fostering collaboration among conservation stakeholders to manage both wild and captive populations as a single conservation unit (Sauve et al., 2022). Combining captive breeding with genomic insights is crucial for mitigating threats (Conde et al., 2011; Farquharson et al., 2021; McGowan et al., 2017), but many species have already lost substantial genetic diversity when these efforts begin (e.g. Dussex et al., 2021; Femerling et al., 2023; Feng et al., 2019; Jackson et al., 2022; Sánchez-Barreiro et al., 2021). Genomic erosion further amplifies drift and inbreeding, driving populations toward an Extinction Vortex (Bosse & van Loon, 2022; Oosterhout et al., 2022; Fagan & Holmes, 2006; Höglund, 2009).

Loss of genetic diversity and increasing realized load significantly challenge long-term conservation efforts (Cavill et al., 2024; Klimova et al., 2022; Lacy, 1997; Oosterhout et al., 2022). Genomic erosion unfolds over generations as population decline creates a ‘drift debt,’ where reduced effective population sizes (Ne) lead to a delayed loss of genetic diversity and an accumulation of inbreeding and deleterious variants that persist across generations (Gargiulo et al., 2024; Pinto et al., 2024). This lag can cause traditional conservation assessments, like the IUCN Red List, to underestimate genetic risks highlighted by genomics (Oosterhout, 2024). While genomic assessments could improve management strategies, comprehensive genetic diversity assessments of pre-bottleneck populations are rare (Cavedon et al., 2023; Speak et al., 2024). Available assessments are often retrospective, relying on pedigree data or limited genetic markers (e.g., Jackson et al., 2022; Russello & Jensen, 2018; Witzenberger & Hochkirch, 2011), which reduces accuracy if pedigrees are incomplete or markers unrepresentative of the whole genome (Allendorf et al., 2010). Whole-genome assessments are essential to understanding inbreeding and genetic load, as bottlenecks elevate realized load by increasing recessive deleterious variants, causing inbreeding depression and fitness loss. (Dussex et al., 2023; Bertorelle et al., 2022). In some cases, purifying selection can purge highly deleterious variants, preserving fitness (Bertorelle et al., 2022; Dussex et al., 2023; Robinson et al., 2023). However, bottlenecks often lead to accumulation of mildly deleterious variants, reducing post-bottleneck fitness (Dussex et al., 2023). Additionally, the benign environment and equalized genetic contribution in captive breeding can reduce the efficacy of natural selection, allowing deleterious variants to persist (Lynch & O’Hely, 2001; Wright et al., 2021).

The whooping crane (*Grus americana*) is a powerful case study for evaluating how conservation strategies can mitigate genetic diversity loss in wild and captive populations. The whooping crane is an iconic, endangered North American bird species, recognized by its distinctive white plumage, bugling call, and intricate courtship dance.The wild population once exceeded 10,000 individuals, but hunting, habitat loss, and disturbance reduced their numbers to around 1,300 by 1870 (Allen, 1952; Canadian Wildlife Service & U.S. Fish and Wildlife Service, 2007). In 1941-1942 the species reached its lowest point with just 16 individuals. By the 1950s, only the US/Canada portion of the North American Central Flyway remained, within a single surviving wild population (Canadian Wildlife Service & U.S. Fish and Wildlife Service, 2007).

The whooping crane gained protection under the U.S. Migratory Bird Treaty Act (1918), the Endangered Species Preservation Act (1966), and the Endangered Species Act (1973), where it was listed as endangered in 1978 (USFWS, 1967, 1970), a status that has been reaffirmed since (CWS, 2011; Canadian Wildlife Service & U.S. Fish and Wildlife Service, 2007). The species has been listed as Endangered in the IUCN Red List since 1994. A captive breeding program began in 1966, and today the captive population includes over 130 individuals across 19 institutions, aiming to retain over 90% of the remaining species’ genetic diversity for the next century through strategic pairings and transfers between institutions (Boardman et al., 2021). Captive breeding minimizes mean kinship and inbreeding, with genetically valuable chicks retained to preserve diversity (Boardman et al., 2021; McAbee & Conkin, 2024). Captive breeding has supported four reintroduction efforts; two have failed and two are ongoing; the Eastern Migratory Population (EMP, est. 2001) and the Louisiana Non-migratory Population (LNMP, est. 2011) (Louisiana Department of Wildlife and Fisheries, 2022; H. L. Thompson et al., 2022). Although reintroduced populations show some natural recruitment, they are not yet demographically self-sustaining and regular releases of captive-reared individuals are used to reinforce and maintain the size of these populations (Caven et al., 2023; Louisiana Department of Wildlife and Fisheries, 2022; H. L. Thompson et al., 2022). The only remnant migratory population, the Aransas-Wood Buffalo Population (AWBP), has naturally grown from 16 individuals in 1941 to around 540 today in protected wintering and breeding grounds without captive supplementation and it is demographically self-sustaining (Caven et al., 2023; McAbee & Conkin, 2024). Currently, there are over 830 whooping cranes globally across remnant, reintroduced, and captive populations (McAbee & Conkin, 2024).

Amid progress toward demographic recovery, the U.S. Fish and Wildlife Service internally discussed downlisting the whooping crane under the ESA, as information was obtained via a Freedom of Information Act request by the Center for Biological Diversity (Caven et al., 2023; Kurose, 2022). Meanwhile, the IUCN aims to reassess the status of all bird species by 2030 to monitor biodiversity progress and guide conservation efforts (IUCN, 2019). The whooping crane’s status under the ESA and IUCN frameworks depends on demographic, range, and threat factors, which should include genetic health (BirdLife International, 2020; Canadian Wildlife Service & U.S. Fish and Wildlife Service, 2007; Caven et al., 2023). Genetic and genomic assessments of extant whooping crane populations have been limited, and population viability analyses are hindered by the lack of recent genomic data (Caven et al., 2023; Miller, 2024; Miller et al., 2021; Traylor-Holzer, 2019). Moreover, insights from conservation genomic studies are inadequately incorporated in extinction risk assessments (Oosterhout, 2024). Existing research indicates that whooping cranes have lost at least two-thirds of their pre-bottleneck mitochondrial DNA haplotypes (Glenn et al., 1999; Jarvi et al., 2001). Captive populations also continue to lose genetic diversity at a modest rate, despite population management best practices (Boardman et al., 2021), which is a likely reflection of the drift debt (Pinto et al. 2024; Jackson et al. 2022).

In this study, we evaluate genomic erosion in the whooping crane using 53 whole genomes spanning the past two centuries across historical, captive, and wild populations. Leveraging a new chromosome-level assembly, we reconstructed demographic history over the last million years and analyzed genetic diversity, inbreeding, and genetic load dynamics for the past 200 years. By estimating both masked load and realized load, we assess the species’ vulnerability to future inbreeding depression. Finally, using the complete pedigree for whooping cranes, we evaluate the impact of captive breeding management on mitigating genomic erosion. With this unprecedented genomic insight, we caution against downlisting the species and advocate for an integrated *in-situ* (wild) and *ex-situ* (captive) management, as proposed in the One Plan approach to conservation.

## Materials and Methods

### De-novo reference genome sequence and assembly

We generated a high-quality reference genome using the Vertebrate Genome Project’s trio v1.6 protocol (Rhie et al., 2021), from two parents and a male offspring collected at the International Crane Foundation in 2019. For each parent we generated Illumina PE 150bp (88X) and used them to sort the haplotypes of Pacbio continuous long reads (CLR; 233X); the sorted reads were used to generate contigs and then scaffolded each haplotype into chromosomes with 10X Genomics (111X), Bionano (417X), Arrima HiC (94X) data. We polished the final maternal and paternal assemblies with the 10X short reads for correcting base call errors and manually curated for structural errors. We chose the maternal haplotype assembly as the reference, as it was the more complete assembly. The offspring’s maternal haplotype assembly has a total length of 1.3 Gb in 39 autosomes, the Z sex chromosomes, 929 scaffolds, and the complete mitochondrial genome. The assembly has a BUSCO (Simão et al., 2015) completeness of 98.4%. The assembly was fully phased and both haplotypes are deposited in NCBI (accession numbers: maternal haplotype: GCA_028858705.1 and paternal haplotype GCA_028858595.1).

### Sample collection, DNA extraction, library preparation and sequencing

#### Modern samples

We collected 37 modern samples (2007-2020): 18 from the ICF (2007-2019) and 19 from wild individuals in the Aransas-Wood Buffalo population (Aransas NWR, TX, USA; WBNP, NWT, Canada; 2009-2020) (Figure 1A, Table S1). ICF, with the largest captive population (36 individuals), maintains a detailed pedigree and fitness data for all birds (S. Liu, 2018; McAbee & Conkin, 2024). Using this information, we classified each captive individual into three different categories depending on how many generations they have been in the captive program (Founders_wildborn, N=6; Captive_early, N=6; and Captive_late, N=6). Founders_wildborn are captive-born individuals whose parental generation were wild individuals (e.g., eggs collected from the AWBP); Early_captive are the first- or second-generation descendants of the founders; and Late_captive are captive-born birds after at least three generations in captivity (Figure 1B).

**Figure 1.**
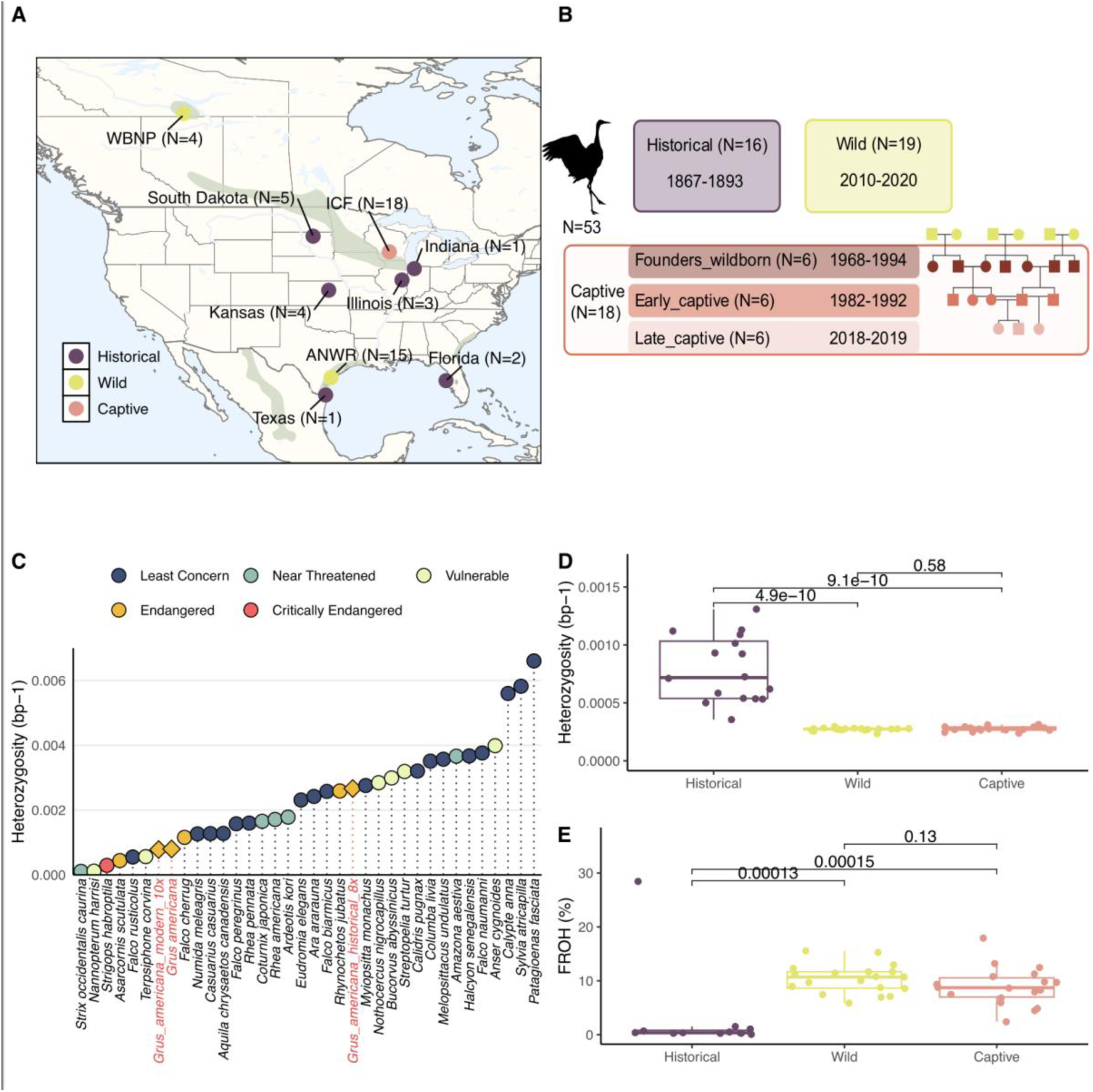
Temporal reduction of genome-wide heterozygosity and increase of inbreeding in whooping cranes. **A)** Map of current distribution of whooping cranes (in green) with the sampling locations of historical, and modern captive (ICF) and wild (WBNP and ANWR) individuals. **B)** Experimental design of historical and modern sampling. Captive individuals include three generations: Founders_wildborn are those whose parental generation were wild individuals, and the eggs were hatched in captivity; Early_captive are the first (or second) generation descendants of Founders_wildborn; and Late_captive are captive-born birds born after at least three generations in captivity. **C)** Genome-wide heterozygosity (all sites) of the whooping crane (red font; *Grus americana*) in comparison with other bird species with different IUCN Red List status. *Grus_americana_modern_10x* represents the average heterozygosity of all modern samples sequenced in this study after downsampling to 10x coverage; *Grus americana* is the reference genome individual at 66.54x coverage; and *Grus_americana_historical_8x* is the historical sample with the highest coverage (MCZ-42511 at coverage 8.37x). Heterozygosity estimates of this historical sample have been corrected with a transition/transversion ratio of 2.89. **D)** Statistically significant difference in genome-wide heterozygosity (only transversions) between historical, wild and captive whooping cranes (Wilcoxon rank-sum test, Historical vs Wild p-value=4.9e-10 and Historical vs Captive p-value 9.1e-10), with a 70% loss in heterozygosity since 1893**. E)** Statistically significant increase of inbreeding (F_ROH_) between historical, wild and captive whooping cranes (Wilcoxon rank-sum test, Historical vs Wild p-value=0.00013 and Historical vs Captive p-value 0.00015), including only samples with coverage >4x.

We extracted DNA from blood stored on FTA cards using the DNEasy Blood & Tissue kit (cat #69506) as described in the protocol for nucleated blood. Briefly, we punched two 2 mm holes from the blood spot on each FTA card using a hole punch. We cleaned the hole punch between samples with 2% NaClO and then with 70% EtOH. We added the punched-out blood spots to a 1.5 mL tube per sample with 20 µL proteinase-K and buffer PBS, bringing the final volume to 220 µL. Finally, we performed the DNA isolation as described in the protocol. We converted the modern extractions into genomic libraries using the NEB Ultra II FS kit (cat #E7805L) with enzymatic shearing. We used 100 ng of DNA as input, shearing extracts for 14 minutes at 37°C, and followed the protocol as described. We size-selected for final library products of 270-370 bp and amplified the libraries for 5 cycles. We pooled the libraries at an equimolar ratio and sequenced the genomic libraries on one lane of 2×150 S4 Illumina NovaSeq 6000 run at Duke University. Modern genomes were sequenced to an average depth of coverage of 13.4x (ranging from 8.1x to 18.3x) (Table S1).

#### Historical samples

We obtained 16 historical toe-pad samples from Kansas University and Harvard Museum of Comparative Zoology collections (Figure 1, Table S1), dated from 1867 to 1893, representing the pre-bottleneck population. We conducted all historic extractions and genomic library preparations in a dedicated clean lab facility at the UCSC Paleogenomics Lab, following protocols described in (Fulton & Shapiro, 2019). We used sterile scalpels to subsample approximately 2-3 mm2 of tissue per extraction, and mechanically homogenized the tissue in 1.5 mL tubes using 1 mL of digestion buffer optimized for hair and tissue (Gilbert et al., 2007). We isolated DNA following (Dabney et al., 2013) and prepared single-stranded Illumina compatible genomic libraries following (Kapp et al., 2021). We amplified the libraries as informed by qPCR results, and sequenced the libraries on an in-house Illumina NextSeq 550 2×75 run to assess sample and library quality. We then pooled the libraries at an equimolar ratio and sequenced the libraries on one lane of 2×150 S4 Illumina NovaSeq 6000 run at Globe Institute Sequencing Core. We increased the sequencing effort for seven of the historical samples to obtain deeper coverage with an extra run in a 2×150 S4 Illumina NovaSeq 6000 at Novogene UK. Historical genomes were sequenced to an average of 4.9x (ranging from 2.9x to 8.4x) (Table S1). Historical samples showed deamination patterns (<10%) typically observed in ancient DNA samples (Figure S2).

### FASTQ trimming, mapping and quality control

We trimmed reads and removed adapters from raw FASTQs with SeqPrep2 (https://github.com/jeizenga/SeqPrep2), retaining reads longer than 30 base pairs and with MQ >20. We aligned the sequences to the maternal haplotype (GCF_028858705.1) using BWA 0.7.17 *mem* (default parameters) for modern samples and *aln* (-l 1024 -n 0.03 -o 2) for historical samples (Li & Durbin, 2009). We removed PCR duplicates with picard MarkDuplicates 2.27.5 (http://broadinstitute.github.io/picard/). We assessed DNA damage for historical samples using mapDamage v2 (Jónsson et al., 2013) and genome-wide coverage with MosDepth v0.3.3 (Pedersen & Quinlan, 2018). For specific analysis as detailed below, we downsampled BAM files to 4x or 10x coverage using samtools v1.10 (Danecek et al., 2021) with *samtools view -s*.

### Genotype calling and filtering

We used snpAD v0.3.1.10 (Prüfer, 2018) to call genotypes in samples with coverage higher than 5x in each autosomal chromosome. Next, we merged individual VCFs per autosome in a single file with bcftools v1.20 (Danecek et al., 2021) *merge* option. We kept only those variants with a read depth between 4 and 50 (included) and a minimum genotype quality of 30 with bcftools *filter*. Finally, we concatenated all VCF files with bcftools *concat*. All the analyses were restricted to the autosomes.

### Demographic history

We used three approaches that apply different inference methods to estimate past demographic history in effective population size (Ne) at different time-scales with the modern samples. For the most recent demographic changes in the past 100 generations we used linkage disequilibrium (LD) estimates with GONE (Santiago et al., 2020), with default parameters and a recombination rate of 3.42 cM/Mb (Cui et al., 2024). We randomised individuals and sites with a Jackknife strategy to obtain a distribution of Ne estimates. For estimates over the past 10,000 years we used site frequency spectrum (SFS) data in StairwayPlot V2 (X. Liu & Fu, 2015, 2020) with a generation time of 13 years (Gil de Weir, 2006), a mutation rate of 1.45e-8 (Zhang et al., 2014), and 8 random breaks. We obtained the folded SFS with ANGSD (-uniqueOnly 1 -remove_bads 1 - only_proper_pairs 1 -C 50 -baq 0 -minMapQ 30 -minQ 20 -doCounts 1 -GL 2 -doSaf 1; Korneliussen et al., 2014). Finally, to obtain the long-term Ne we used Pairwise Sequentially Markovian Coalescent (PSMC) (Li & Durbin, 2011) with the reference genome sample mapped to itself, as it has the highest coverage (66.54x). We ran *psmc* with the following parameters: -N30 -t5 -r5 -p “4+30*2+4+6+10” following (Nadachowska-Brzyska et al., 2015) and performed 100 independent bootstrap runs.

### Relatedness

We used ANGSD (Korneliussen et al., 2014) to obtain the genotype likelihoods in binary format (-doGlf3 -GL 2) separately in the modern and historical datasets with the following filters: -uniqueOnly 1 -remove_bads 1 -noTrans 1 -only_proper_pairs 1 -C 50 -skipTriallelic 1 -minMapQ 30 -doMajorMinor 1 -doMaf 1 -SNP_pval 1e-6. With the resulting files we ran NgsRelate v2 (Hanghøj et al., 2019) to obtain the theta kinship coefficient. Standard kinship distribution values were used to classify individuals as unrelated when their kinship coefficient is 0, third-degree or higher relatives for values between 0 and 0.0625, second-degree for values between 0.0625 and 0.1875, and first-degree for values between 0.1875 and 0.375.

### Sample pedigree and inbreeding coefficients

We used the R package FamAgg (Rainer et al., 2016) to construct a pedigree of 1,836 captive-bred individuals from the Studbook (S. Liu, 2018). We calculated inbreeding coefficients for all individuals using the function inbreeding from the ribd package (Vigeland, 2020), and calculated the relatedness of sampled individuals using the kinship function of the FamAgg package, which calculates relatedness as the chance that a locus is identical between the two individuals. Here, in the absence of inbreeding, a parent and child would have a calculated kinship of 0.25, and a selfing cross equates to a kinship of 0.5.

### Heterozygosity

We estimated heterozygosity per sample using ANGSD (Korneliussen et al., 2014) and winSFS (M. S. Rasmussen et al., 2022). We first extracted a list of high-quality sites by estimating genotype likelihoods with the following parameters: -uniqueOnly 1 - remove_bads 1 -only_proper_pairs 1 -rmTrans 1 -C 50 -minMapQ 20 -minQ 20 -setMinDepth 3 -setMaxDepth 50 -doCounts 1 -doMajorMinor 1 -GL 2 -doGlf 2 -doMaf 2. Next, we ran ANGSD with the list of high-quality sites to obtain their site allele frequencies (-doSaf 1). Finally, we ran winSFS to obtain the site frequency spectrum (SFS) per sample.

For the comparison between historic and modern samples we removed transitions to account for DNA damage and, to test for the effect of different depths of coverage in our samples, we also calculated heterozygosity in BAM files downsampled to 4x.

For comparisons within modern samples, we estimated heterozygosity at all positions (including transitions) using BAM files downsampled to 10x with ANGSD using the following parameters: -uniqueOnly 1 -remove_bads 1 -only_proper_pairs 1 -noTrans 0 -C 50 -baq 0 -minMapQ 30 -minQ 20 -setMinDepth 3 -setMaxDepth 50 -doCounts 1 - GL 2 -doSaf 1. Then we employed winSFS as before.

Next, we evaluated how much genome-wide genetic diversity the whooping crane has in comparison to other bird species with a range of conservation statuses (Table S2). For each species, we mapped raw sequencing reads to the reference genome with bwa 0.7.17 (Li & Durbin, 2009) for Illumina sequenced genomes or pbmm2 v1.13.1 for genomes sequenced with Pacbio (https://github.com/PacificBiosciences/pbmm2) using default settings. Next we estimated genome-wide heterozygosity from the BAM files by first obtaining genotype likelihoods with ANGSD (Korneliussen et al., 2014) with the following parameters: -uniqueOnly 1 -remove_bads 1 -only_proper_pairs 1 -C 50 -baq 0 -minMapQ 30 -minQ 20 -setMinDepth $minDP -setMaxDepth $maxDP -doCounts 1 - nThreads 10 -GL 2 -doSaf 1. MinDP and MaxDP were set to ⅓ and 2 times the average coverage in each genome. Next, SFS was estimated with realSFS -fold 1. We included the heterozygosity estimate of the historic sample with highest depth of coverage in the multi-species comparison. To account for the different estimate obtained when retaining only transversion sites, we corrected the estimate using the transition/transversion ratio of 2.89, which we estimated from the modern samples.

### Inbreeding with Runs of Homozygosity

We estimated runs of Homozygosity (ROH) using ROHan (Renaud et al., 2019) on samples with coverage >4x. For historical samples, we precalculated damage patterns by running the script estimateDamage.pl (on the 50 bp at 5’ and 3’ of reads) provided by the software. We used these estimates to identify ROH with a minimum size of 1Mb on the autosomes with the following parameters: –size 1000000 --rohmu 2e-5 --deam5p ${sample}.5p.prof --deam3p ${sample}.3p.prof --auto autosomes.txt.

For modern samples, we estimated ROH using the same parameters as above but without the damage pattern profile files. Given the differences in coverage between historical and modern samples, we also downsampled modern samples to 4x and re-ran ROHan. When restricting the analysis to only modern samples, we downsampled them to 10x.

We estimated inbreeding age by converting ROH lengths (cM) to generation with the formula G = 100/(2*cM) (E. A. Thompson, 2013), using a 3.42 cM/Mb recombination rate (Cui et al., 2024) and a 13-year generation time (Gil de Weir, 2006).

### ROHbin (Runs of Homozygosity per bin)

We used a custom approach, ROHbin (***R****uns **O**f **H**omozygosity per **bin***), to determine the private ROHs variation (as determined by low heterozygosity regions) for each wild and captive population. The method partitions the genome into bins of 1 Mb and leverages a statistical framework to estimate bins that contain unique variation in each of the tested groups. For each bin, ROHbin uses an empirical Bayes moderated F-statistic with the R function *ebayes* from the limma package (Ritchie et al., 2015) in R v. 4.1 (R Core Team, 2022) to determine statistical significance and effect size (as logFC) for each comparison. We considered bins with a p-value < 0.05 to be significantly different between groups and likely to contain private variation in the wild or captive population.

### Genetic load

#### SnpEff

We polarized the discovered variants into ancestral and derived using four outgroups: the three closest ancestral nodes to the whooping crane (Figure S1) from the B10K genome alignment of 363 avian species (Feng et al., 2020) and the reference genome of one sister species (*Grus nigricollis* GCA_004360235) (Zhou et al., 2019). We obtained the ancestral nodes fasta sequence in the hal file (Feng et al., 2019) using hal2fasta (Hickey et al., 2013). We fragmented each fasta sequence into 150bp long sequences using bedtools windowMaker v2.30.0 (Quinlan & Hall, 2010) and subsequently mapped them to our whooping crane reference genome with bwa mem v0.7.17 (Li & Durbin, 2009). We obtained variants using bcftools mpileup and bcftools call 1.15.1 (Danecek et al., 2021), and combined them to the whooping crane VCF with bcftools merge. We filtered the resulting VCF to remove fixed positions and keep variants with a genotyping rate of 40%. We also restricted the analysis to those variants with a 100% of genotyping rate across the four outgroups. We annotated putative deleterious variants with SnpEff (Cingolani et al., 2012).

From the annotated VCF we extracted those variants classified as High, Moderate, and Low impact. High-impact variants are assumed to have a high (disruptive) impact in the protein, probably causing protein truncation, loss of function (LoF) or triggering nonsense-mediated decay (i.e., stop codons, splice donor variant and splice acceptor, start codon lost, etc.). Moderate impact variants are non-disruptive variants that might change protein effectiveness (i.e., missense variants). Low-impact variants are mostly harmless or unlikely to change protein behavior (i.e., synonymous variants). We considered only variants with a minimum read depth of 5. For the analysis including modern and historical samples we excluded variants only present in the modern population because our power to uncover variants in the post-bottlenecked population is much greater than the discovery power we have for the more diverse, historical, ancestral population.

To approximate the genome-wide genetic load, we counted derived alleles relative to the outgroup ancestral alleles as putatively deleterious. We separated the total genetic load into heterozygous load and homozygous load by counting the number of derived alleles with low, moderate, and high predicted levels for homozygous alleles (multiplied by two) and heterozygous alleles. The homozygous counts are a proxy of the realized load (deleterious variants that express fitness effects), and heterozygous counts are a proxy of the masked load (deleterious variants whose fitness effects remain largely hidden). Heterozygous variants can partially express their deleterious fitness effects (Bertorelle et al., 2022), but in the absence of dominance coefficients (h) estimates, we assumed that h∼0 for moderately and highly deleterious mutations (Charlesworth & Willis, 2009).

To account for the effect of sequencing depth, we downsampled two modern samples to 5, 10 and 15x coverage (EB31_S103 and EB33_S105). Given that differences in depth of coverage between historical and modern samples can create biased allelic counts, we normalized the Moderate and High-impact allele counts by Low-impact allele counts, effectively correcting for variant discovery power.

Next, we estimated the historical to modern frequency change of deleterious variants (low, moderate and high). For each category of variants, we estimated the observed allele frequency per site in each population (modern or historical). We first tested if the observed frequency in each category had a significant chance over time with a binomial test. Then we calculated 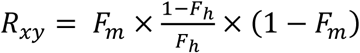 (Do et al., 2015), normalizing the R_xy_ values for both high and moderate categories with the low category, as previously done for allelic counts. We performed a jackknife analysis to estimate the variance around the R_xy_ values. Finally, we estimated the proportion of homozygous load found inside and outside runs of homozygosity (ROH) using the bedtoolsr R package (Patwardhan et al., 2019). The counts of homozygous load were normalized by the total homozygous sites inside and outside ROHS.

#### LoadLift

Next, we further assessed genetic load in the captive-bred and wild populations using the LoadLift pipeline (Speak et al., 2024) to obtain CADD scores derived from model species (chicken). This analysis is complementary in different ways. First, contrary to SnpEff we retained CADD scores within evolutionary conserved regions of the genome (ultra conserved elements, UCEs) as mutations in these regions are more likely to be unconditionally deleterious (i.e., independent of their genetic and environmental background). Also, since CADD scores are continuous, this method allows for a conversion of predicted deleterious scores into selection coefficients to estimate the genetic load components, i.e., the realized load and masked load (Bertorelle et al., 2022). We limited the inference to the 0.72% portion of the genome contained in UCEs and their 2000 bp flanking regions using the Phyluce pipeline (Faircloth, 2016). We retained sites within the UCE with CADD scores present in all 37 modern individuals, where the chicken reference genome and whooping crane sample data contained the same alleles and where sites were not fixed across all 37 samples (144/536774 sites retained). CADD scores are PHRED scaled and therefore not additive, and hence, we converted the CADD scores to rank values. We must note, however, that this linear scale of rank-scores is only a crude proxy of the complex and largely unknown distribution of selection coefficients, which we know is non-linear. Hence, comparing the load components within and between individuals needs to be interpreted with caution.

The genetic load, realized load and masked load were calculated for every locus following the formulas in Bertorelle et al., (2022): 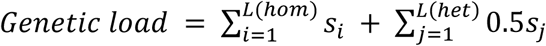; 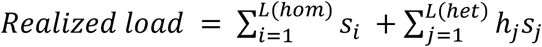; 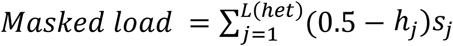. The rank score of selection coefficient (s_i_) was inferred from their CADD score. The dominance coefficient (h_j_) was categorized in function of the CADD score; for variants with CADD<10 we assumed h=0.3, for 10<CADD<20 h=0.15, for 20<CADD<30 h=0.02, and CADD>30 h=0. We thus assumed there is a negative relationship between s and h and that more highly deleterious mutations are expected to be more recessive (Charlesworth & Willis, 2009).

To investigate how the genetic load varies as a function of kinship (taken from the pedigree) and relatedness (estimated from genomic data), we analyzed the resulting genetic load component values of offspring from hypothetical wild-wild, captive-captive and captive-wild crosses, assuming Mendelian segregation ratios of the parental alleles (Speak et al. 2024).

## Results

### Temporal patterns of genetic diversity, inbreeding and genetic load

We estimated the genome-wide diversity of the 2012 captive-born whooping crane reference genome individual (mapped short-reads to 66.54x coverage). The obtained value of 0.0008 het x bp^-1^ is on the low end of the diversity range observed across 34 bird species from diverse phylogenetic lineages (Figure 1C). We obtained the same value when the sample was downsampled to 10x coverage, highlighting that this depth is sufficient to confidently estimate genome-wide metrics. The genome-wide diversity of whooping cranes before the historical population size minimum (sample MCZ-42511 collected in 1893, with 8.37x coverage) is three times higher, 0.00267 het x bp-1 (Figure 1C). This represents a statistically significant 70% loss of heterozygosity through time (Figure 1D). To avoid any potential bias related to difference in sequencing coverage (Figure S3), we estimated heterozygosity after downsampling the entire dataset to 4x and found the same pattern (Figure S4). The loss of heterozygosity is coupled with a marked increase of inbreeding over time as evidenced by the increased fraction of the genome in runs of homozygosity (FROH) (Figure 1E). Historical individuals showed no evidence of inbreeding as on average only 0.5% of their genome was contained in ROH (0.08-1.5%, with one outlier at 28.4%), while modern individuals consistently showed high FROH with an average of 9.6% of their genome in ROH (2.4-17.9%), with 3.91% of the genome in ROHs longer than 10Mb consistent with recent inbreeding (Figure 1E, Figure S5 and Table S1)

Genetic load in modern populations significantly decreased for moderate and high categories (Figure 2A), driven by a loss of heterozygous load (Figure 1D). The homozygous load did not increase significantly, possibly because the heterozygous variants were more likely to be purged than drift to fixation. To avoid biases, we restricted the above analysis to sites present in at least one historical sample and excluded private modern variation. We normalized the Moderate and High-impact derived allele counts by the Low impact derived allele counts to mitigate the impact of differences in sequence coverage between modern and historic datasets (Figure S6).

**Figure 2.**
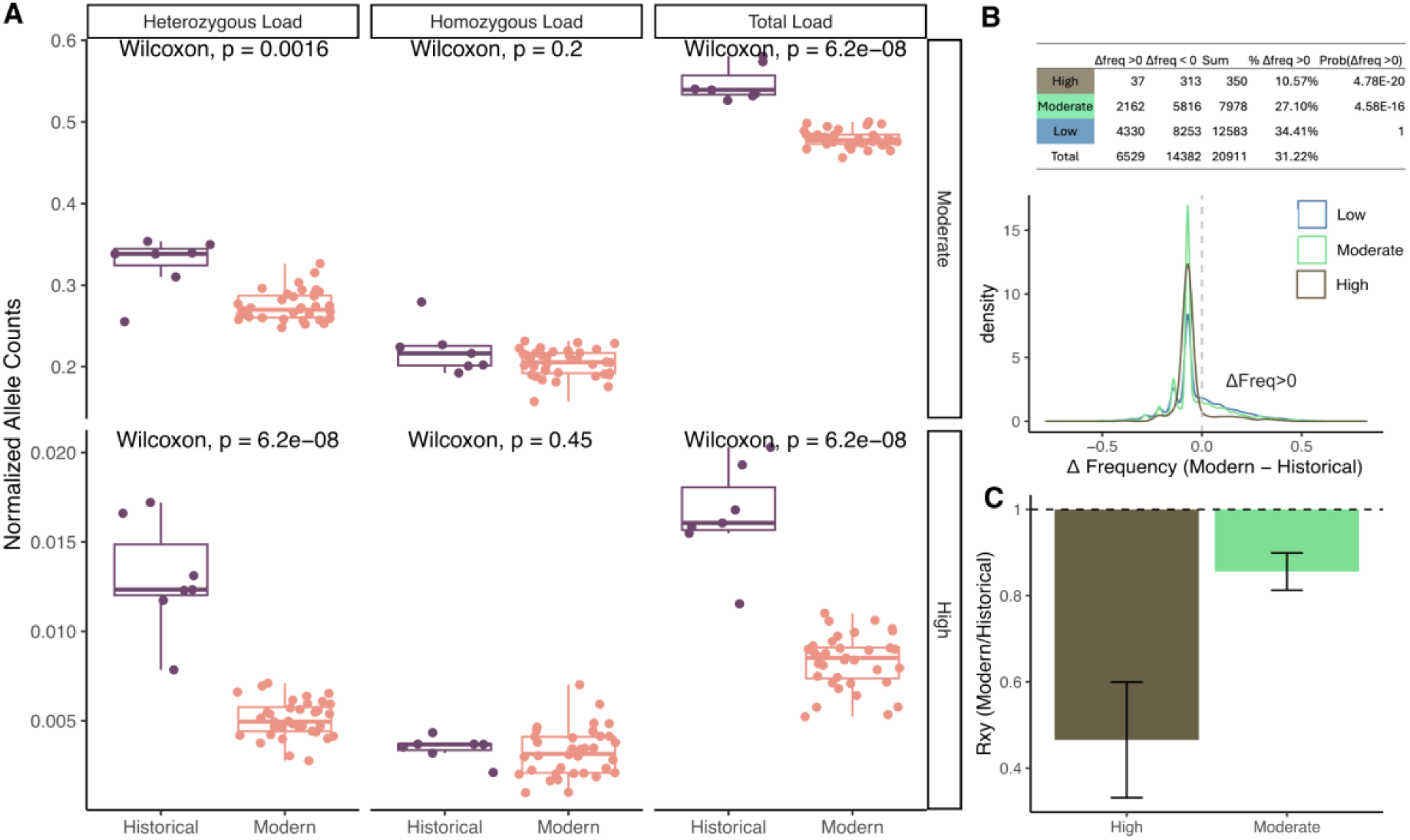
Temporal dynamics of genetic load. **A)** Normalized derived allele count of deleterious variation (moderate and high impact) at heterozygous, homozygous and total load. **B)** Variant counts that increased and decreased in frequency from historical to modern with a binomial statistical test; and distribution of Δfrequency differences (Modern-Historical) of deleterious variants (low, moderate and high). **C)** *Rxy* ratio of derived alleles between modern and historical samples for the High and Moderate impact deleterious variants (normalized by the Low *Rxy*). Normalized *Rxy*<1 indicated relative frequency deficit of the corresponding category in the modern samples compared to the historical samples. Error bars represent the variance after jackknife resampling of variant sites.

Next, we observed a lower frequency of high-impact variants (average DeltaFreq = - 0.053) compared to moderate (average DeltaFreq = −0.030) and low-impact variants (average DeltaFreq = −0.017) in the modern time point. High-impact variants also had a significantly smaller proportion with DeltaFreq > 0 than the other categories (Figure 2B). Additionally, only 10.48% of high-impact variants increased in frequency in modern samples without reaching fixation, compared to an increase in frequency in modern samples of 26.52% for moderate and 33.46% for low-impact variants. The proportion of variants increasing versus decreasing in frequency is statistically significant for both high-impact (p = 4.78E-20) and moderate-impact (p = 4.58E-16) categories (binomial test). Together, these results suggest that selection is operating against highly deleterious variation, keeping the frequency of those variants at low levels (i.e. purging). To further test whether the purging of deleterious variants occurred during the extreme bottleneck, we calculated the ratio of derived alleles (Rxy) between modern and historical samples for each impact category. We found a significant depletion of highly deleterious variants (Figure 2C). Overall, the genetic load dynamics are consistent with an effect of genetic drift and purifying selection against deleterious variation, where most deleterious alleles were likely lost at random after the bottleneck, and some (mostly highly deleterious) were purged by negative selection.

### Demographic history of whooping cranes

Our PSMC reconstruction indicates that the whooping crane population has been relatively small for a long time (Figure 3A), with a harmonic mean Ne of 11,444 (between 5,000 and 1×10^6^ years ago) and a maximum Ne of 23,515 (∼80,000 years ago during the Late Pleistocene). After this point, our reconstruction from more recent time with StairwayPlot shows a steady decline that continues through the Holocene (Figure 3B) until recent times. A concordant signature picked up from reconstruction of the most recent past with GONE, showing that the most recent bottleneck started around 300 years ago (mid 17th century) and continued until 100 years ago (Figure 3C). At this point (8 generations ago, around 1916), we estimated the lowest Ne = 37.

**Figure 3.**
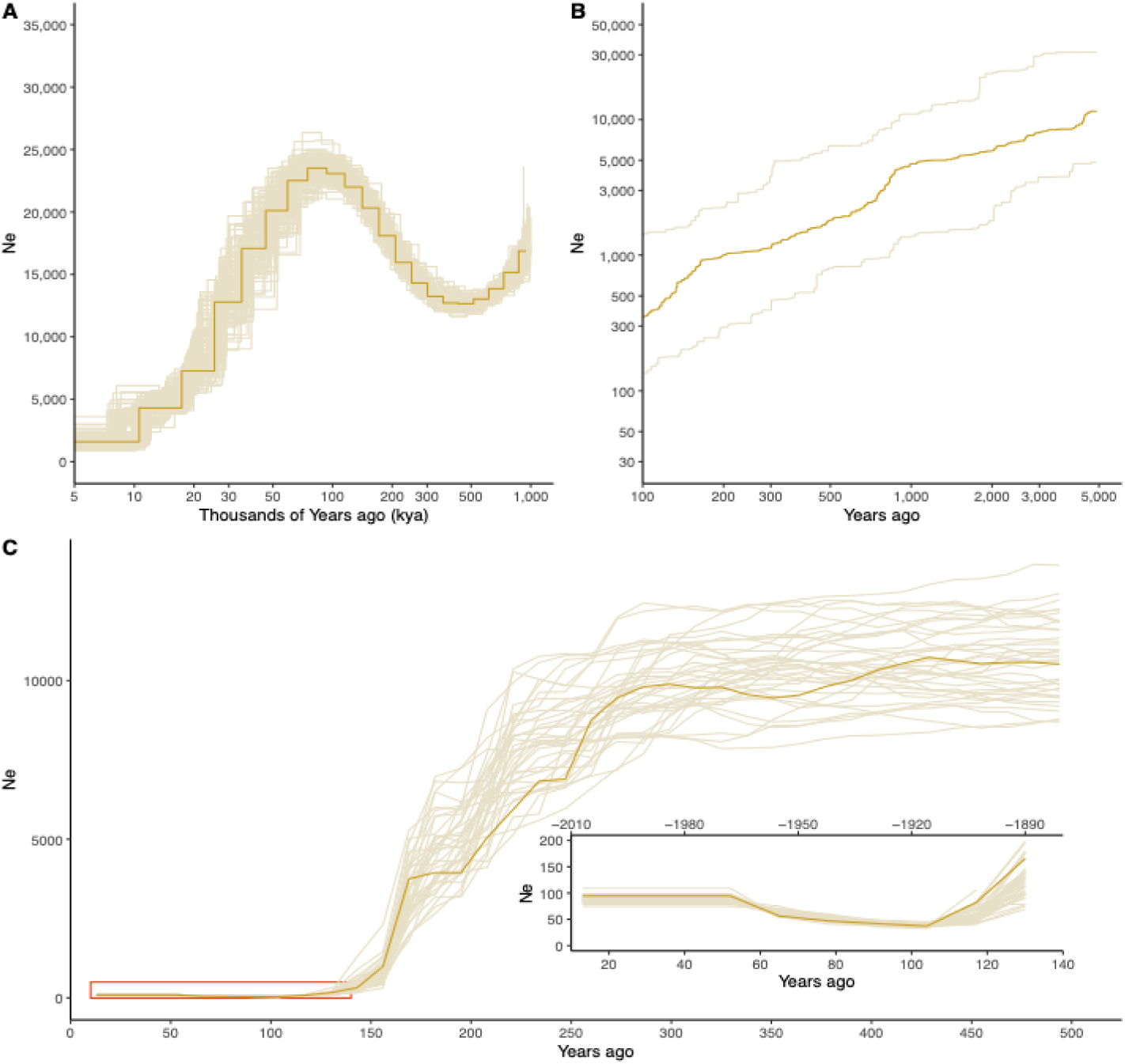
Demographic trajectory of the whooping crane through time. **A)** PSMC analysis of the reference genome sample. **B)** StairwayPlot of all modern samples and **C)** GONE analysis using all modern samples in the last 500 years (or ∼40 generations). The red square highlights the last 140 years as shown in the inserted plot at the bottom left.

### Effect of captive management in whooping cranes

We integrated the pedigree information of the captive population and classified individuals according to their generation in captivity (Figure 1B and Table S1). The average pairwise kinship coefficient is 0.0095 (sd = 0.03), well below the third-degree relatedness threshold (0.0625) (all estimates in Table S3). The estimated kinship within “Founders_wildborn” individuals was relatively low, but increased in subsequent generations, within the “Early_captive” cohort, to then be reduced again among the “Late_Captive” (Figure S7), evidence that the captive population was likely founded with unrelated individuals and that captive breeding management has actively reduced relatedness.

We detected a non-significant trend of loss of heterozygosity across captive generations (Figure 4A). Moreover, wild and “Late_Captive” individuals, which represent a similar generation after the bottleneck, show similar levels of genome-wide heterozygosity (Figure 4A). Inbreeding accumulated after the bottleneck in the captive breeding program, but also in the wild unmanaged population (Founders_wildborn=7.94%, Early_captive=8.43%, Late_captive=10.31% and Wild=10.36%) (Figure 4B; Table S1 and Figure S5). We used a dataset downsampled to 10x to avoid potential biases due to differences in coverage (Figure S8) and we detected no difference in FROH using all data available or downsampling to 10x (Figure S9).

**Figure 4.**
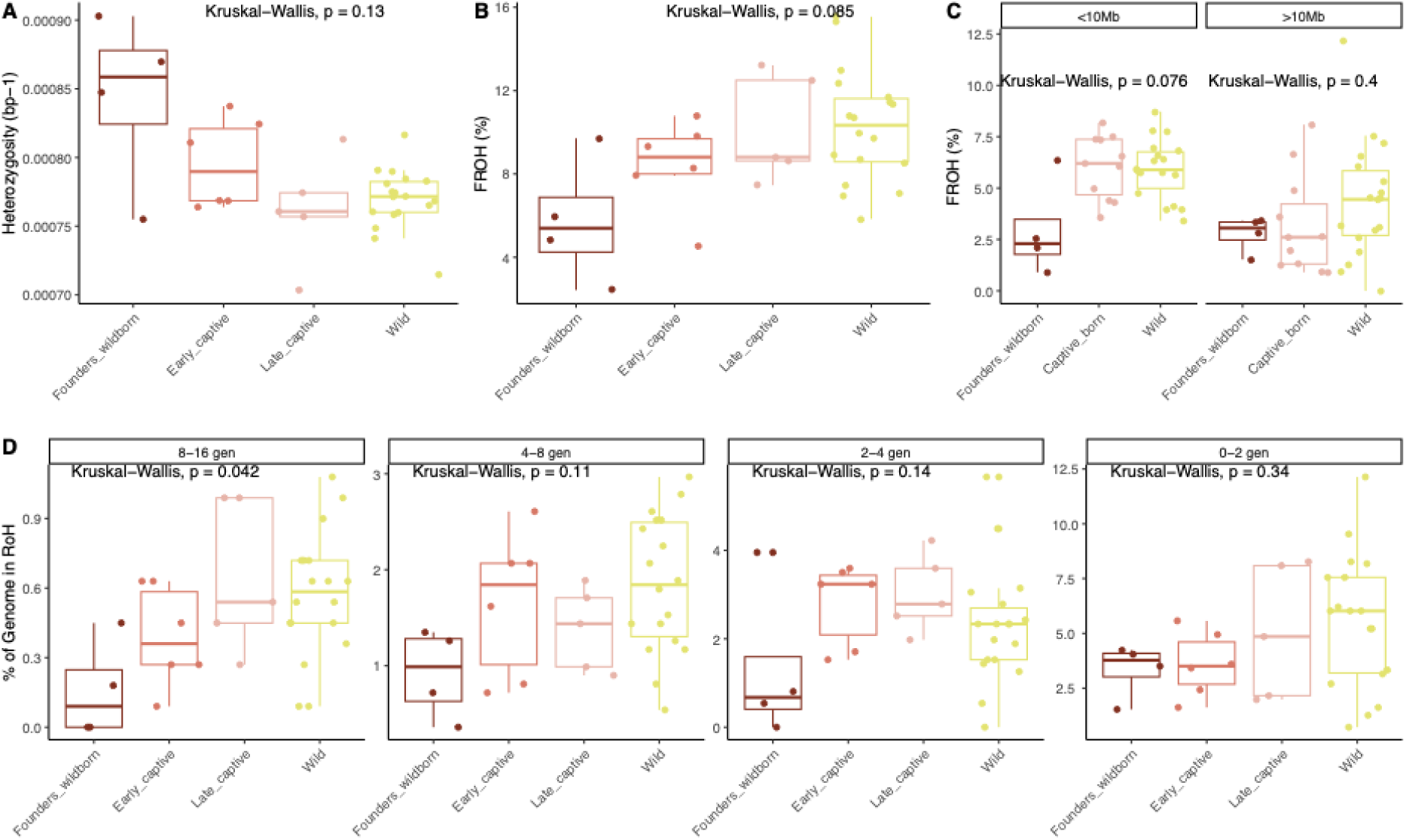
Genomic diversity and inbreeding in a captive time series and wild individuals. **A)** Genome-wide heterozygosity. **B)** Inbreeding calculated as F_ROH_. **C)** Inbreeding (F_ROH_) grouped by size: smaller or larger than 10 Mb and merging early and late captive-bred birds (Captive_born). **D)** Inbreeding (F_ROH_) after stratifying ROHs by coalescence time (generations ago). In all plots, global statistical significance is calculated with a Kuskal-Wallis test.

The coalescence time of most ROHs is within the last 4 generations or 50 years (Figure S10). Interestingly, the observed accumulation of FROH since the bottleneck is driven by ROHs smaller than 10Mb (Figure 4C, Figure S11), reflecting background inbreeding due to historical demographic events. In line with this evidence, we only identify a statistically significant increase of the proportion of the genome in ROHs for the oldest generation bin (8-16 generations ago, approx. 100-200 years ago) (Figure 4D). On the other hand, long ROHs have not accumulated between the first and last generation of captivity (Figure 4C). This may indicate that while captive breeding practices avoid inbreeding, the accumulation of background inbreeding was unavoidable in the absence of genomic data and due to the small founding captive population size (N=35).

We did not detect any changes for the genetic load across successive captive generations or in the wild population (Figure S12). However, we identified a statistically significant accumulation of genetic load inside ROHs compared to outside ROHs in the Late_captive and wild populations (Figure S13). We observed a significant reduction of the genetic load for low and moderate categories outside ROHs (in the Late_captive population compared to the wild (Figure S14). Overall, this pattern suggests that captivity might be reducing the efficacy of selection, but further analyses decomposing different ROH classes and mutation score predictions with a larger sample size is required to be conclusive.

The predicted deleterious scores from both SnpEff and CADD were consistent in ranking mutation severity. SnpEff-classified deleterious mutations had higher average CADD scores than low-impact mutations (moderate: 20.7, high: 23.5, low: 2.60; Wilcoxon rank sum, p < 2.2e-16) (Figure 5A). For sites categorized by SnpEff as moderately deleterious, derived alleles (both heterozygous and homozygous) had significantly lower average CADD scores compared to homozygous wild-type alleles across all modern samples (Wilcoxon rank sum, p < 2.2e-16). Furthermore, no homozygous alleles were detected for mutations classified as highly deleterious, aligning with the expected effects of strong purifying selection (Figure 5B).

**Figure 5.**
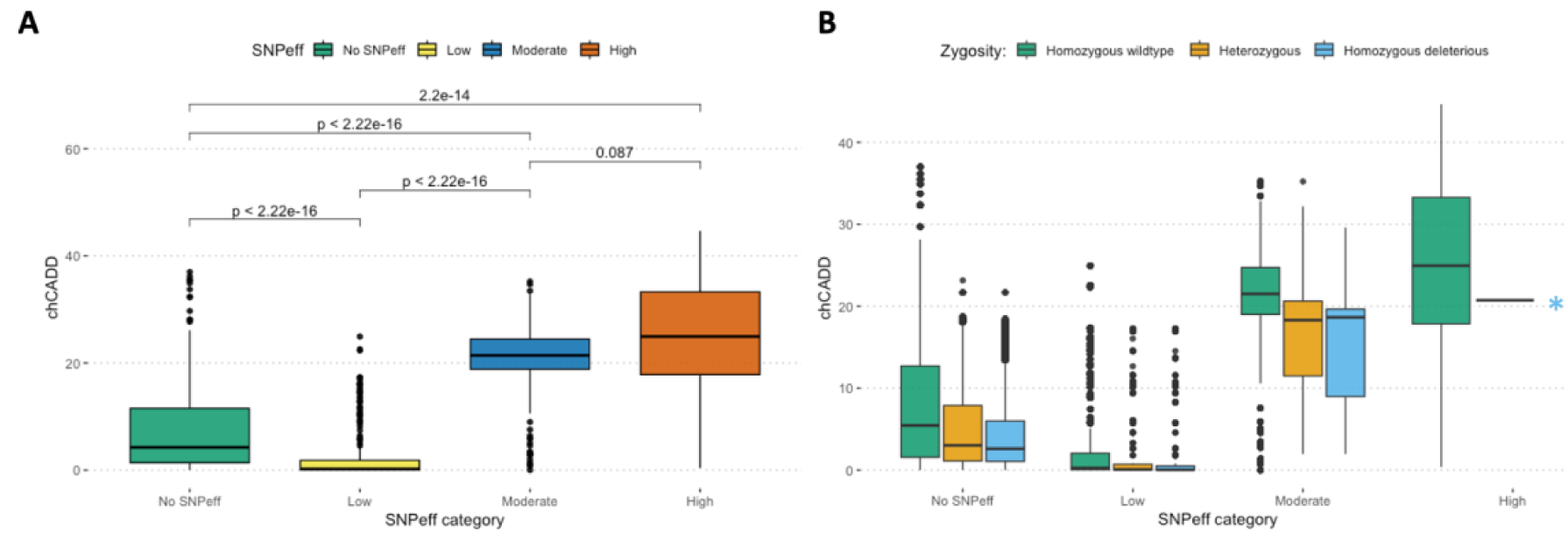
A comparison of lifted chCADD scores and SnpEff categories across 37 whooping crane samples. **A)** One-to-one comparison of SnpEff categories and chCADD scores. **B)** Zygosity compared across the different SnpEff categories, with individuals homozygous for the wildtype chicken reference allele (non-scoring) (Green), heterozygous for the deleterious allele with a chCADD score (Orange) and homozygous for the deleterious allele with a chCADD score (Blue). A star (*) is used to denote that no sites categorised as highly deleterious by SnpEff were homozygous for the deleterious allele.

Converted chCADD scores to genetic load estimates (see Methods) revealed that the realized load exceeded the masked load in whooping cranes (Figure 6A), likely reflecting the effects of prolonged inbreeding and genetic drift from sustained small population sizes and serial bottlenecks. Inbreeding depression can be caused by a higher expression of genetic load in more inbreed individuals, however, we detected no significant correlation between genetic load components and FROH (genetic load: R² = 0.0012, p = 0.8376; realized load: R² = 0.0111, p = 0.5342; masked load: R² = 0.0111, p = 0.5349). We proffer two explanations. First, ROHs contain mostly historical tracts already purged of highly deleterious mutations (Bosse & van Loon, 2022). Second, due to the low level of masked load, inbreeding does significantly elevate the realized load, which is consistent with our findings reported in Figure 2A (i.e., homozygous load does not increase over time).

**Figure 6.**
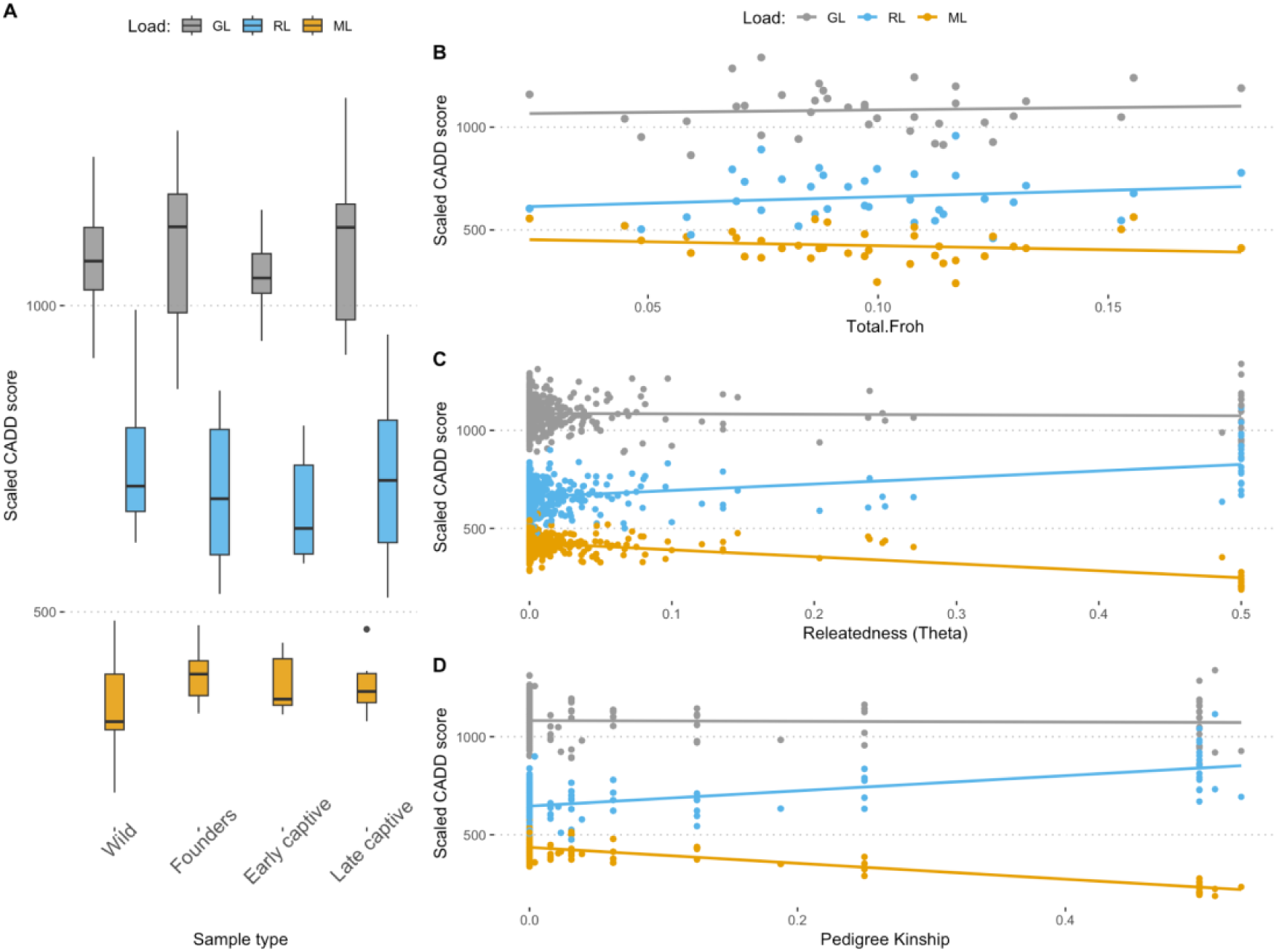
Genetic load components within ultra conserved elements (UCE) estimated from chicken CADD (chCADD) scores. **A)** Individual genetic load was calculated for sites within the UCE the realized load (RL) (Orange), masked load (ML) (Blue) and the genetic load (GL) (Grey). **B)** The GL, RL and ML of each of the 37 modern individuals as a function of their FROH, **C)** the GL, RL and ML of potential crosses compared to the genetic relatedness of the parents, and **D)** the GL, RL and ML of potential offspring of crosses as a function of the relatedness of their parents based on the pedigree.

We used converted chCADD scores to predict the genetic load in offspring from hypothetical matings, assuming Mendelian segregation ratios. Results revealed that strategic pairings could slightly minimize expressed genetic load, as the predicted realized load and masked load were significantly affected by parental relatedness (realized load: R² = 0.2388, p = 2.272e-08; masked load: R² = 0.6277, p < 2.2e-16) (Figure 6C). Offspring from closely related pairs exhibited higher realized load, as masked load in the parents was converted into realized load in inbred offspring (Figure 6A). However, the rate of increase in realized load was modest compared to other heavily bottlenecked species, the pink pigeon (Speak et al., 2024), suggesting that most of the whooping crane’s masked load has already been converted, rendering the species relatively robust to future inbreeding. Similar patterns emerged when using pedigree-based relatedness values (genetic load: R² = 0.0035, p = 0.5872; realized load: R² = 0.2559, p = 6.75e-07; masked load: R² = 0.6711, p < 2.2e-16) (Figure 6D). These findings indicate that complete pedigree records in the captive program have effectively managed inbreeding and stabilized genetic load. Overall, while contemporary inbreeding may slightly increase realized load, this effect is likely to remain minor due to the current low masked load in the captive and wild populations.

### Complementary genetic diversity between wild and captive populations

Analyzing 1 Mb genomic bins, we found that 5.1% of the wild genomes (∼57 Mb) had significantly lower heterozygosity than the captive genomes (Figure 7A, Table S4), while 3.2% of the captive genomes (∼35 Mb) showed lower diversity than the wild genomes (Figure 7B). These regions likely contain private genetic variation specific to each group. Excluding founders no longer contributing genetic diversity, the amount of private variation in the captive individuals (Late_captive) reduced to 1.16% (Table S4). These results align with the accumulation of FROH and ongoing diversity loss in captive generations (Figure 4B), though this analysis is limited by only six samples, reducing statistical power. All analyses yielded consistent results and patterns using smaller bin sizes of 500 kb or 250 kb (Table S4).

**Figure 7.**
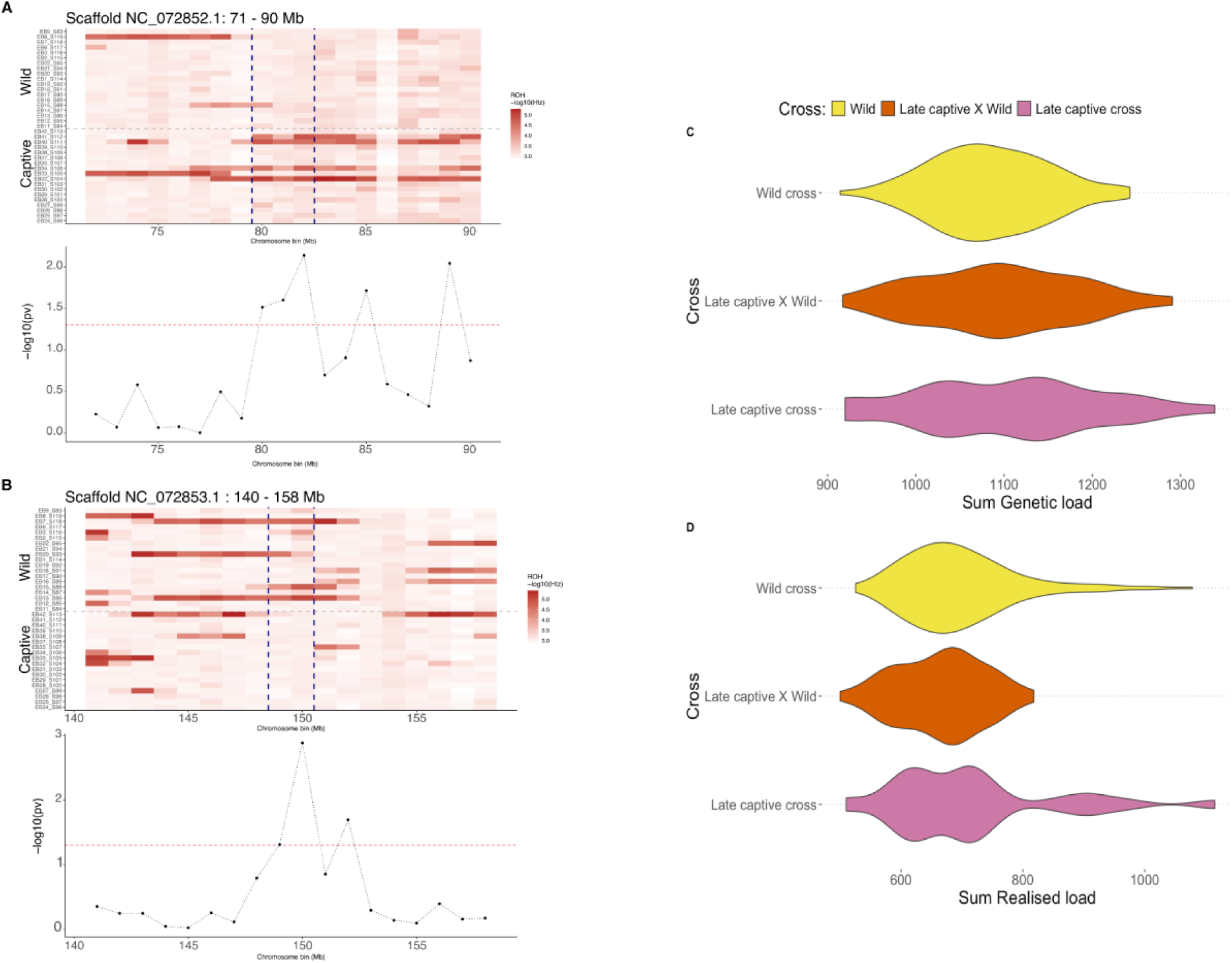
Complementarity of genetic diversity and genetic load between captive and wild populations. **A)** Example of a region with significant enrichment of low heterozygosity in the captive population. ROHbin identifies regions private to each group, showing trends in low heterozygosity. **B)** Example region with significant enrichment of ROH, private to wild populations. In panels A-B, rows represent individuals, columns show 1Mb genome bins, and color intensity reflects heterozygosity (-log10 scale). **C)** Distribution of genetic load across potential crossings. **D)** Realized load from crosses between wild-wild (yellow), wild-captive (orange), and captive-captive (pink) individuals

Finally, we evaluated the genetic load in hypothetical optimal pairings to test whether captive-wild crosses could reduce the realized load. The genetic load was similar for wild-wild, captive-captive, and captive-wild crosses (Figure 7C). However, genomics-informed selection of captive-wild crosses could produce offspring with the lowest realized load by masking deleterious alleles (Figure 7D). There was no significant difference between average realised load of captive-wild crosses and either wild-wild or captive-captive crosses (Kruskal-Wallis rank sum test, df = 2, p = 0.6738, *x*^2^ = 0.79). However, optimal captive-wild crosses could reduce realized load by 44.44% compared to the worst captive-captive pairings, and by 43.22% compared to the worst wild-wild pairings. Similarly, the optimal captive-wild cross could reduce realized load by 26.14% when compared to the mean of the five best crosses from all of the sampled individuals if selected based on pedigree kinship alone.

## Discussion

The whooping crane faced extreme bottlenecks, nearly leading to extinction. Conservation actions, including habitat preservation, hunting restrictions, and breeding programs, have since enabled population recovery to over 830 individuals, sparking debate on potential downlisting from Endangered to Threatened under the ESA and also their eventual potential downlisting in the IUCN Red List (Caven et al., 2023).

In this study, we provide temporal genomic evidence of the severe consequences of the whooping crane past population collapse. While we demonstrate the success of the captive breeding program in keeping recent inbreeding rates low, we also caution against downlisting the species due to the significant sustained genomic erosion. Contemporary whooping crane populations have lost approximately 70% of their ancestral genome-wide diversity since 1893. Reduced heterozygosity could limit the population’s ability to adapt to environmental changes (Femerling et al., 2023; Hoelzel et al., 2019; Hoffmann et al., 2017), and increased Inbreeding, evidenced by elevated FROH values, further exacerbates this issue by increasing the expression of deleterious alleles (i.e., realized load) potentially reducing population viability (Bertorelle et al., 2022). While most highly deleterious variants have likely been purged, remaining deleterious mutations still contribute significantly to a prevalent realized load through recent time and across all populations. Increasing gene flow between captive and wild populations could help reduce the realized load and introduce novel genetic variation, potentially enhancing their long-term viability.

### Evolutionary dynamics of genomic erosion through time

As previously described in other temporal genomic datasets from heavily bottlenecked species (e.g., Seychelles magpie-robin - Cavill et al., 2024; Kākāpō - Dussex et al., 2021; Seychelles paradise flycatcher - Femerling et al., 2023; Crested ibis - Feng et al., 2019; Pink pigeon - Jackson et al., 2022; Cotton-top tamarin - L. Rasmussen et al., 2023; Black rhino - Sánchez-Barreiro et al., 2023; Eastern gorilla - van der Valk et al., 2019), whooping cranes show a strong (70%) reduction in genome-wide diversity over time and marked inbreeding accumulation (Figure 1). Long-term Ne estimates indicate the whooping crane declined over the past 1,000 years. This trajectory aligns with the late-Quaternary extinction (Bergman et al., 2023; Koch & Barnosky, 2006; Svenning et al., 2024), which saw the extinction of two European crane species (*Grus primigenia and Grus melitensis*; Northcote, 1984; Northcote & Mourer-Chauviré, 1988), a fate narrowly avoided by the whooping crane.

Still, the whooping crane population crashed even further in the last 300 years (Figure 3C), overlapping with European settlement of the Americas in areas where the whooping crane nested. Historical records show the population size dropped to around 16 individuals in the 1940s, and our recent genomic Ne estimates are similar (Ne = 37 around 1916). The temporal analysis of genome-wide heterozygosity loss offers greater confidence in Ne estimates. A loss of 70% since 1894 or 9.2 generations is equivalent to the level of genomic erosion expected in a population with Ne = 3.8, underscoring the fragile genetic health of the population. The Ne estimates at similar time points differ markedly across the three demographic reconstruction methods—likely due to their reliance on different data sources (Nadachowska-Brzyska et al., 2022). Nevertheless, their trends are highly consistent. This allows us to confidently infer that: (1) the ancestral whooping crane population was never large, (2) it has been in steady decline for the last 100,000 years, and (3) the population was already small during the two bottlenecks in the mid-1800s and mid-1940s.

The whooping crane’s demographic history of long-term small Ne and its recent abrupt collapse provide important context for understanding the dynamics of genetic load over time. We found no change in homozygous genetic load between historical and modern samples, possibly due to our focus on variants shared across historical samples, which may slightly bias against variants unique to modern samples. The rationale behind this strategy is linked to the small size of our historical dataset (7 historical samples passing the coverage filter, compared to 37 modern samples). Without this filtering step, the variant discovery rate would be skewed towards modern samples, falsely suggesting the presence of novel mutations when they may simply be more detectable in the larger, higher-quality modern dataset.

In line with theoretical expectations, we observed a reduction in the genetic load, driven by a decrease in deleterious heterozygous alleles. The significant decrease observed in high-impact deleterious alleles suggests that natural selection removed highly deleterious variants but not moderate impact variants during the prolonged and severe bottleneck. This pattern mirrors what has been observed in other heavily bottlenecked species, such as the Alpine Ibex (Grossen et al., 2020), suggesting that purging of highly deleterious variants may have enhanced the species’ resilience to inbreeding depression. However, some deleterious mutations increased in frequency and could become fixed in future generations due to ongoing inbreeding or genetic drift.

### Captive breeding genomics

A common challenge for breeding programs is that they are often established with a small founding stock of assumed unrelated individuals (Willoughby et al., 2017). The whooping crane captive population was founded with 35 individuals (Mirande et al., 1991). Here, we sampled six of these founders and found that pairwise kinship values among them were lower than expected for a third-degree relationship. In the next generation in captivity, relatedness increased, probably due to a still very small captive population size. But in further generations this value went down, as the breeding program avoided pairing closely related individuals using the rich pedigree information in this species.

Although the captive breeding program has avoided crosses between highly related individuals, loss of heterozygosity and accumulation of ROHs have continued across generations. This highlights the long-term genetic impact of founder events, where small initial populations continue to affect diversity despite inbreeding management (Boakes et al., 2007). Notably, the accumulated inbreeding is mostly due to short ROHs, suggesting the captive program effectively prevents long ROHs from recent inbreeding, but can do little to avoid background (historical) inbreeding. These findings emphasize the importance of maintaining a well-documented and complete pedigree, but also highlight that genomic data is crucial to supplement pedigrees, allowing to account for cryptic relatedness and historical inbreeding. Our analysis of *in-silico* optimal mate pairings illustrates this by showing that reduced kinship crosses could help maintain a lower realized load.

In contrast to our CADD analysis, where the realized load exceeded the masked load, our SnpEff results show that the homozygous allele counts are lower than heterozygous counts. Different filtering schemes might have contributed to this discrepancy. Furthermore, our CADD analysis assumes a range of dominance coefficients for the estimating the load components within ultra conserved elements (UCEs), and thus is not directly comparable to the genome-wide SnpEff estimates. The frequency, selection and dominance coefficients of variants at UCEs may differ from those elsewhere in the genome. For example, an analysis of 29,938 polymorphisms within 2189 UCEs in the human genome showed that the polymorphism density within UCEs (one per 22.7 bp) was higher than the genomic average (one per 21.0 bp). However, the frequency of the derived allele was extremely low (less than 6% occur at a frequency of >1%) (Habic et al., 2019). This underscores the need to better understand how *in-silico* genetic load predictions translate into fitness effects in endangered species.

### One Plan Approach to *in-situ* and *ex-situ* management

The One Plan Approach aims to integrate in-situ and ex-situ conservation efforts by involving all stakeholders (Sauve et al., 2022; Speak et al., 2024). We identified a modest amount of private variation in captive and wild populations, suggesting that combining genetic pools could preserve some genetic diversity and reduce inbreeding. Our genomics-informed management (Speak et al., 2024) shows that selecting optimal pairs could reduce realized genetic load by 20.78% compared to random mating. Selecting the best crosses from between the captive and wild individuals could help to reduce the realised load, theoretically improving fitness and population viability. Optimizing crosses may have only a modest effect, as the whooping crane’s realized load exceeds its masked load, suggesting that most masked load has already been converted to realized load, providing some resilience against future inbreeding depression.

One method to achieve genetic exchange is through egg collection from wild breeding sites, as cranes typically raise only one chick from two eggs (Bergeson et al., 2001). Studies suggest collecting one egg does not impact nesting success (Boyce et al., 2005), though some argue it may reduce recruitment in years with favourable breeding conditions (Cannon et al., 2001). With minimal impact, genetic material could be transferred by collecting one egg from a two-egg nest in the remnant population, transporting it with an incubator, and rearing it in captivity for release into reintroduced populations (Kuyt, 1996). The same positive effect could be achieved by supplementing the captive population. The Canadian Wildlife Service and partners have explored collecting a small number of eggs at the remnant wild population (AWBP) to bolster captive genetic diversity (Compass Resource Management Ltd, 2023). Thus, our results suggest these concerted *in situ* and *ex situ* actions could offer genetic benefits to the reintroduced and captive populations.

Translocation efforts must be carefully managed. While increasing genetic diversity is essential, gene flow from the remnant population could inadvertently introduce masked load, potentially leading to higher realized load after future inbreeding. Further studies with larger sample sizes and fitness data could help clarify the genetic variation across captive and wild populations. For instance, gene flow from wild populations could counteract adaptations to captivity that may have accumulated in the captive population (Frankham, 2008). Additionally, ecological and behavioral factors are crucial, as reintroduced populations (EMP & LNMP) exhibit higher adult mortality and lower recruitment than the remnant wild population (AWBP) (Louisiana Department of Wildlife and Fisheries, 2022; H. L. Thompson et al., 2022; Wilson et al., 2016). Given the success and larger size of the AWBP, genetic introductions from the captive population may be unnecessary. However, AWBP genetic variation could benefit captive and reintroduced populations that are not yet self-sustaining (LDWF, 2022; Thompson et al., 2022). For any population management actions intended to improve population genetics and conservation, outcomes should be closely monitored to determine effectiveness (Gitzen et al., 2016).

### Translating genomic insights into effective conservation management

Our genomic analysis underscores the lasting impact of severe bottlenecks on whooping cranes and highlights the importance of genomically informed captive breeding to maintain genetic health and resilience. Despite demographic recovery through *in-situ* and *ex-situ* conservation action, our results indicate persistent effects of reduced Ne and increased genomic erosion within the wild and captive populations. Given these findings, we advise against downlisting the whooping crane from Endangered to Threatened under the ESA and from Endangered to Vulnerable under the IUCN Red List. We instead, recommend maintaining and expanding current conservation actions. Furthermore, our findings underscore the necessity of integrating genomic data into both *in-situ* and *ex-situ* conservation efforts.

Our study also highlights the importance of sampling and biobanking populations before population bottlenecks take hold, to establish a genetic baseline and contextualise genomic estimates for future predictions and conservation actions (Cavill et al., 2024; Díez-del-Molino et al., 2018; Femerling et al., 2023; Jensen et al., 2022). Genomic tools, as used here, offer a comprehensive framework for identifying inbreeding depression, genetic load, and adaptive potential, supporting unified conservation strategies to improve species survival (van Oosterhout et al., 2022; Segelbacher et al., 2022). By working with the International Crane Foundation and utilizing these tools, we are taking critical steps toward data-driven decisions that enhance genetic diversity, reduce inbreeding risks, and improve long-term recovery outcomes for the whooping crane.

## Acknowledgements

The authors would like to thank the University of Kansas Natural History Museum and The Harvard Museum of Natural History for providing samples for this study. Kim Boardman for providing the whooping crane Studbook. We would like to thank the Whooping Crane Tracking Partnership, which has included the U.S. Fish and Wildlife Service, U.S. Geological Survey, Canadian Wildlife Service, Crane Trust, and Platte River Recovery Implementation Program. Particularly, we want to thank D. Brandt, F. Chavez-Ramirez, along with many volunteers, interns and staff of cooperating institutions and contractors for safely capturing wild whooping cranes. We also thank the ICF Crane Conservation and Conservation Medicine department staff for excellent care and coordinated management of the captive cranes used in this study. This work was funded by the European Research Council (StG ERODE, 101078303 and CoG Extinction Genomics 681396); and the Danish National Research Foundation (Center for Evolutionary Hologenomics, DNRF143). Further support was obtained from NERC ARIES PhD studentship (T209447), the Chester Zoo Conservation Scholar and Fellow Scheme, a Research Training Support Grant (RTSG; 100162318RA1) and by The Howard Hughes Medical Institute.

## Data availability statement

All raw data generated in this study has been uploaded to the European Nucleotide Archive (ENA; Accession no.: PRJEB80530). The metadata of these samples can be found in Supplementary Table S1. Code for data analysis can be accessed through GibHub https://github.com/claudefa/WhoopingCrane_genomics, https://github.com/pollicipes/ROHbin https://github.com/saspeak/LoadLift.

## Author Contributions

Design of the study: HEM Sample Collection by: HEM, BKH

Resources by: HEM, MTPG, BS, EDJ Laboratory work by: MCJ, GF, JB

Data analysis: CF, HEM, SS, GF, CP, XW, JAR, BM, JC, YS, LA

Data interpretation: CF, HEM, AJC, SS, CvO, JAR, OF, EDJ Writing: CF, HEM, AJC

Editing and final approval of the manuscript: All Authors

