## Supplementary Figures for "Persistent genomic erosion in whooping cranes despite demographic recovery"

### Supplementary Tables

Supplementary tables can be found in the excel file: SupplementaryTables.xlsx.

#### Table S1

Metadata of all sequenced whooping crane samples, including their origin, location, year of collection, coverage, studbook information, heterozygosity and inbreeding ($F_{ROH}$).

#### Table S2

Heterozygosity of a set of bird species with different IUCN red list status with their accession number and coverage.

#### Table S3

Relatedness estimated as theta (kinship coefficient) between each pair of captive and wild (modern) whooping cranes.

### Supplementary Figures

#### Figure S1

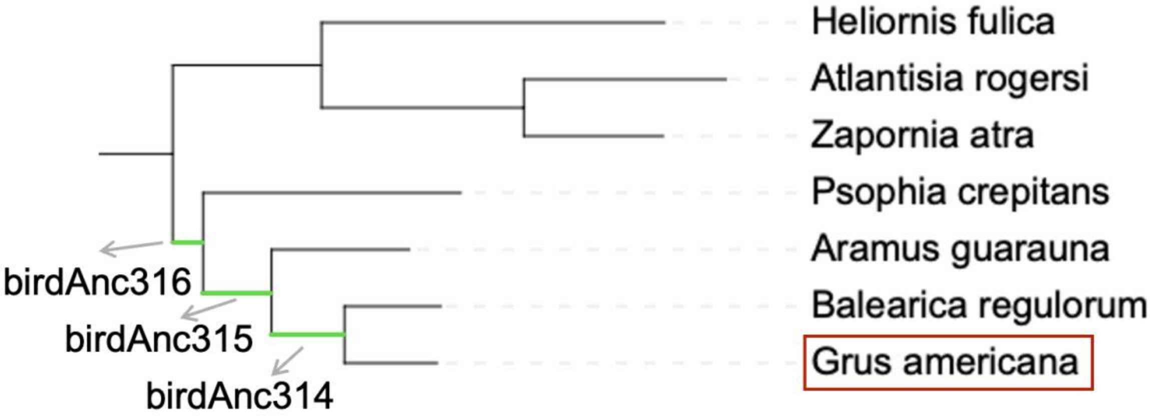

**Figure S1 | Phylogenetic positioning of the whooping crane (*Grus americana*)** from (Feng et al., 2020) to highlight the selected nodes for the polarization of ancestral and derived alleles in the whooping crane.

#### Figure S2

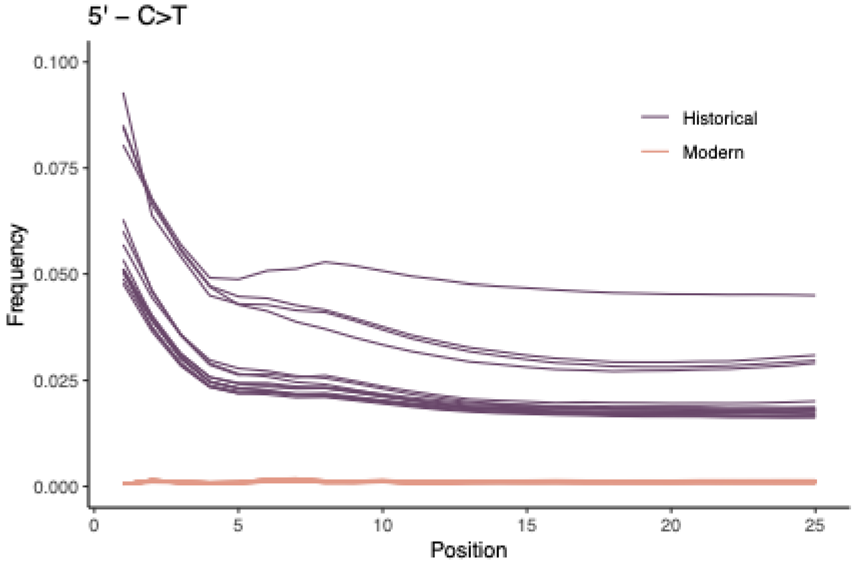

**Figure S2 | Damage patterns (C>T substitution) at 5' of reads**, for historical and modern samples.

### Figure S3

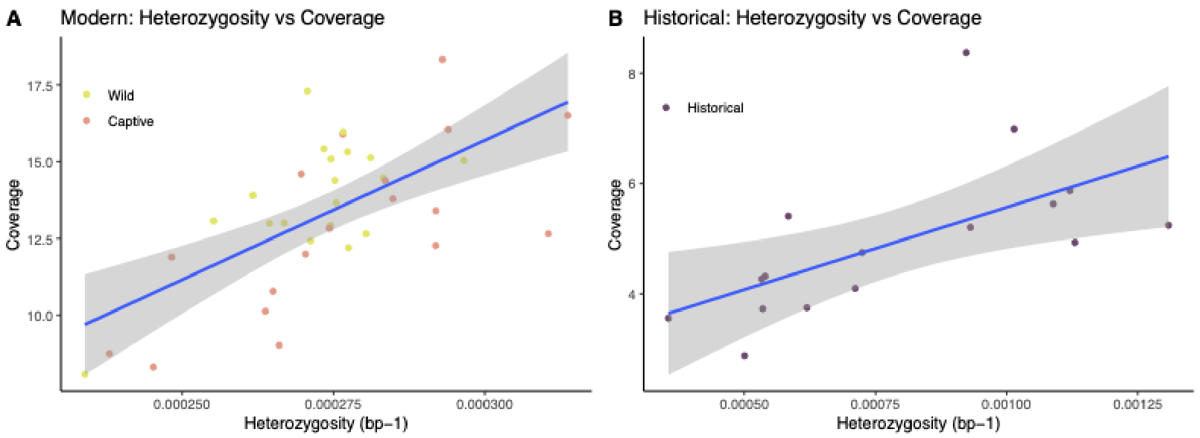

**Figure S3 | Correlation between depth of coverage and heterozygosity (only transversions)** in **A)** modern (adjusted R-squared: 0.3935; p-value 1.953e-05) and **B)** historical samples (adjusted R-squared: 0.3396; p-value: 0.01051).

### Figure S4

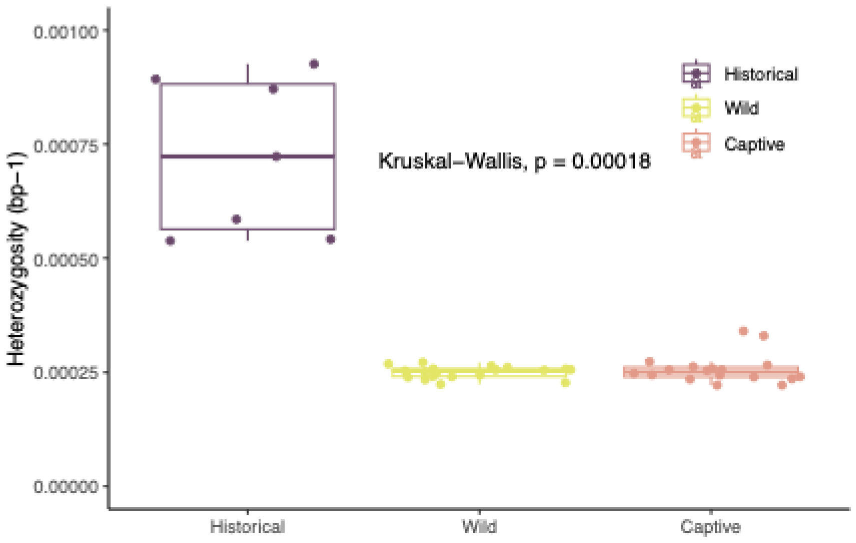

**Figure S4 | Genome-wide heterozygosity (only transversions) after downsampling bam files to an average coverage of 4x.** There is a statistically significant loss of genome-wide heterozygosity between historical, wild and captive whooping cranes (Kruskal-Wallis, p-value=0.00018)**.**

### Figure S5

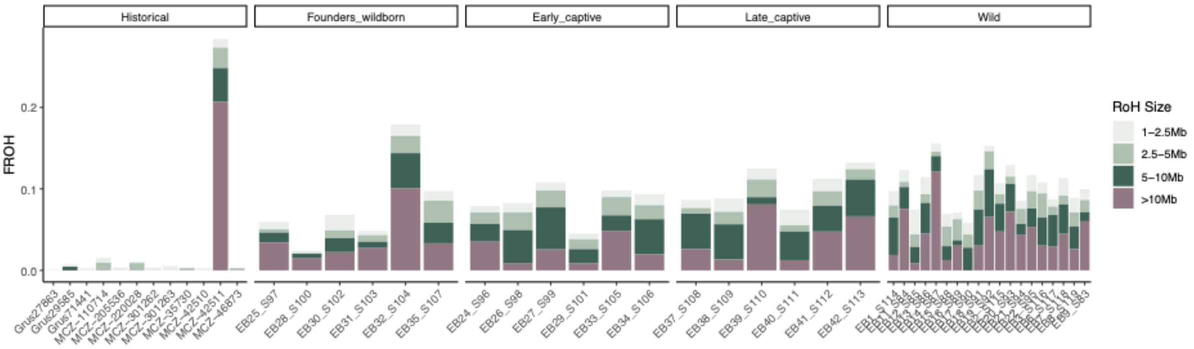

**Figure S5 | Proportion of the genome in Runs of Homozygosity ($F_{ROH}$)** in samples with at least 4x coverage.

### Figure S6

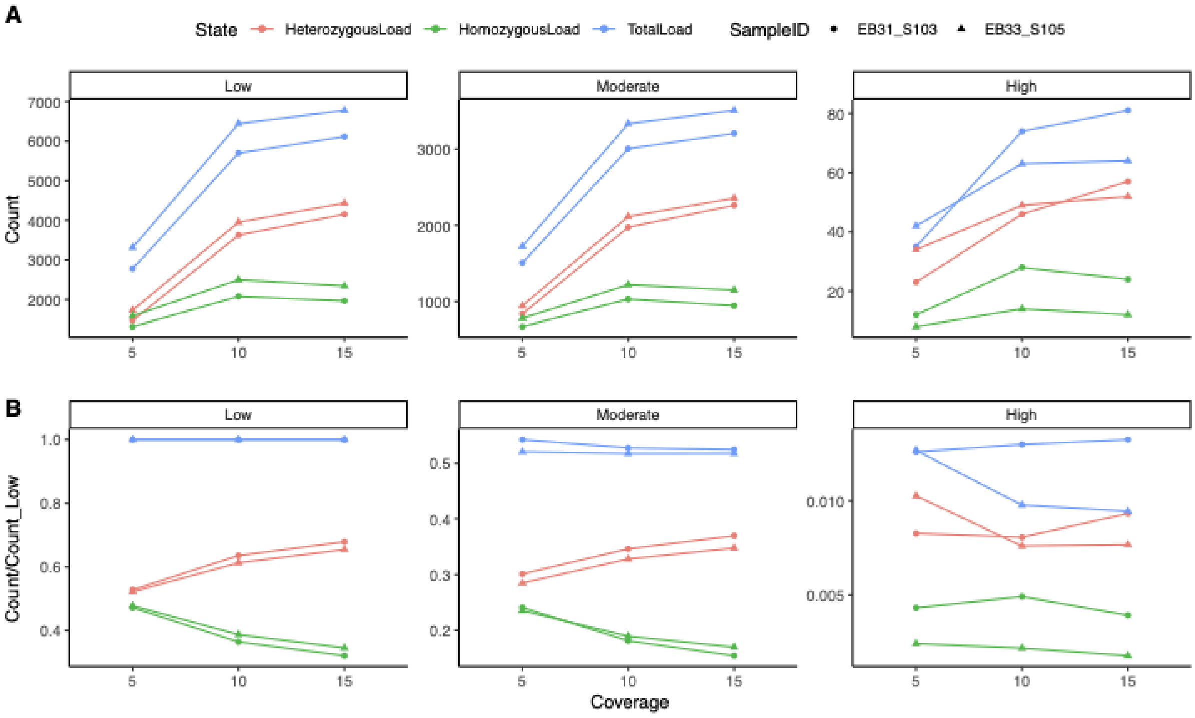

**Figure S6 | Effect of coverage in the detection of deleterious variants in two samples. A)** At 5x coverage the capacity to detect deleterious variants is impaired, especially for heterozygous load. Between 10x and 15x the difference is negligible. **B)** After normalization with total low allele count, the effect of low coverage is mitigated although there is still a tendency to overestimate homozygous load while underestimating heterozygous load.

### Figure S7

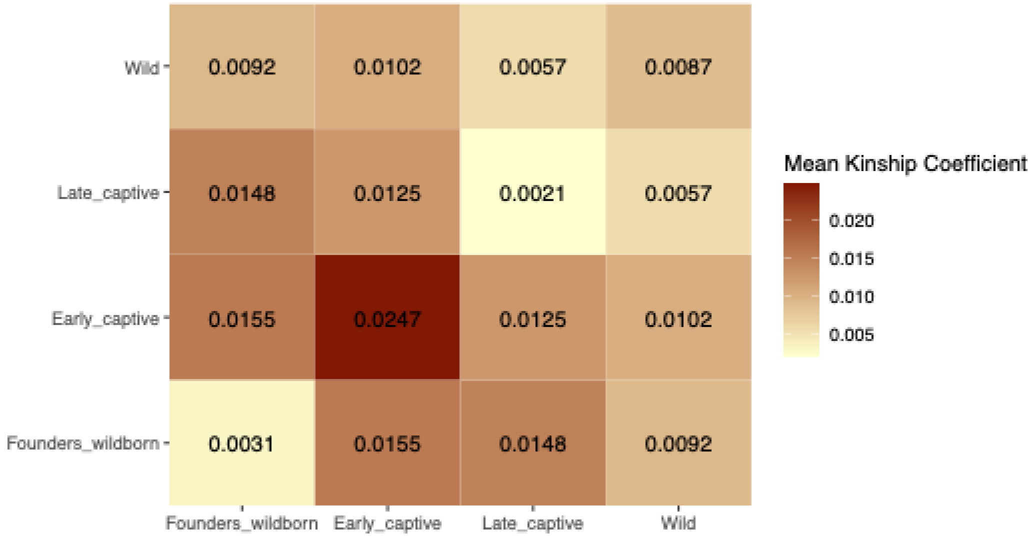

**Figure S7 | Average kinship coefficient (theta)** between and within captive and wild whooping cranes.

### Figure S8

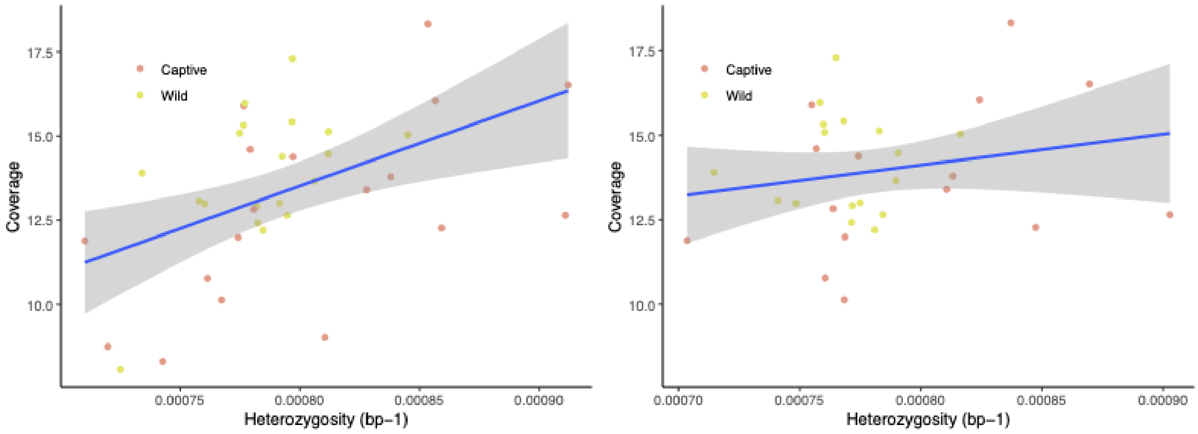

**Figure S8 | Correlation between depth of coverage and heterozygosity (all positions) for modern samples** in **A)** all dataset (adjusted R-squared: 0.2068, p-value: 0.002748) and **B)** after downsampling to 10x coverage and excluding samples with coverage <10x (adjusted R-squared: 0.01021, p-value: 0.2576).

### Figure S9

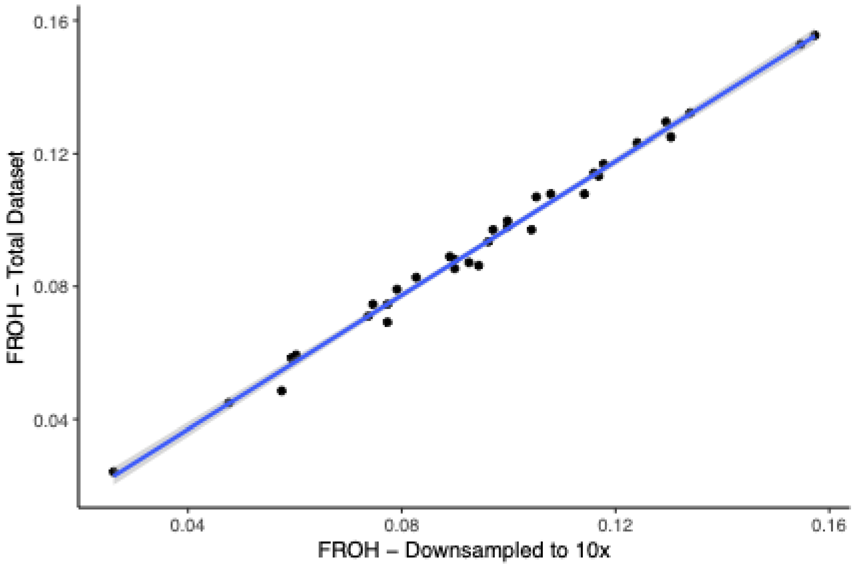

**Figure S9 | Comparison of $F_{ROH}$** between the complete dataset and after downsampling to 10x (adjusted R-squared: 0.991, p-value: < 2.2e-16).

### Figure S10

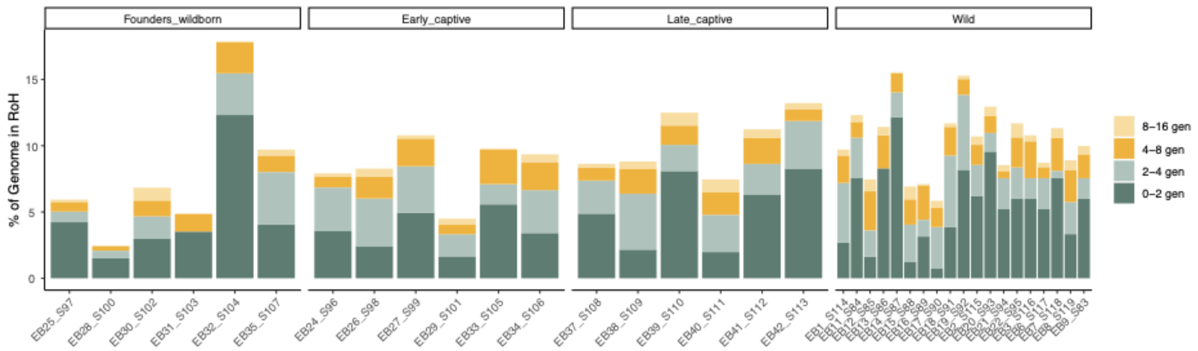

**Figure S10 | Percentage of the genome in Runs of Homozygosity ($F_{ROH}$) stratified by their coalescence time** in modern samples with at least 4x coverage. We obtained the age of ROH in generations using a recombination rate of 3.42 cM/Mb and a generation time of 13 years.

### Figure S11

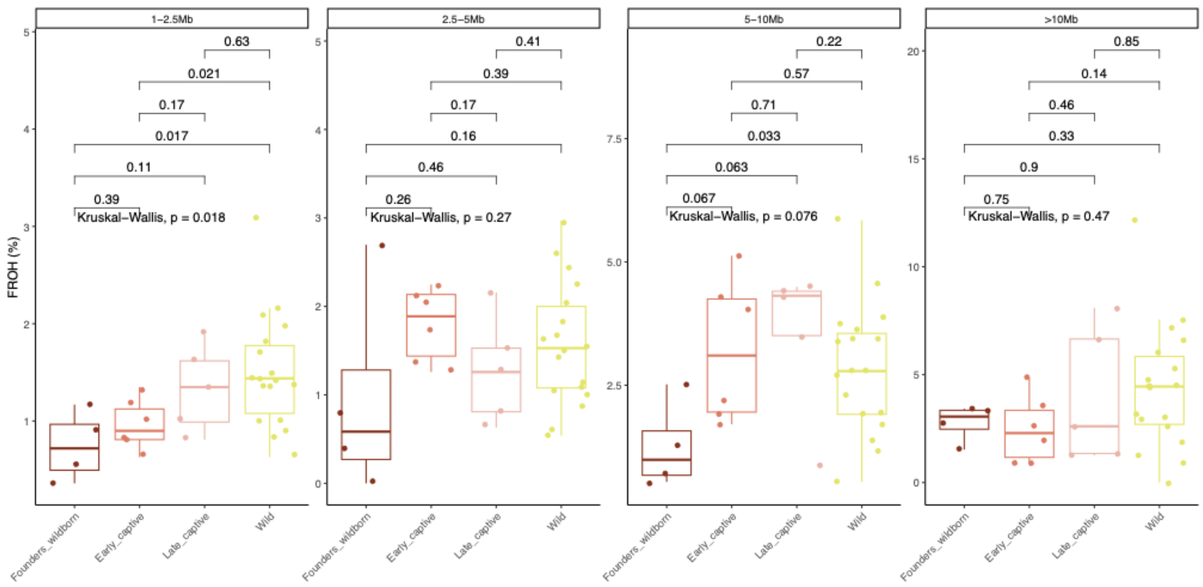

**Figure S11 | Accumulation of runs of homozygosity in different generations in captivity and in the wild.** FROH is segregated by ROH size: 1-2Mb, 2-5Mb, 5-10Mb and >10Mb. Statistical testing in pairwise comparisons are done with the Wilcoxon Rank Sum test, while the overall comparison with a Kruskal-Wallis test. This data only includes samples with at least 10x coverage.

### Figure S12

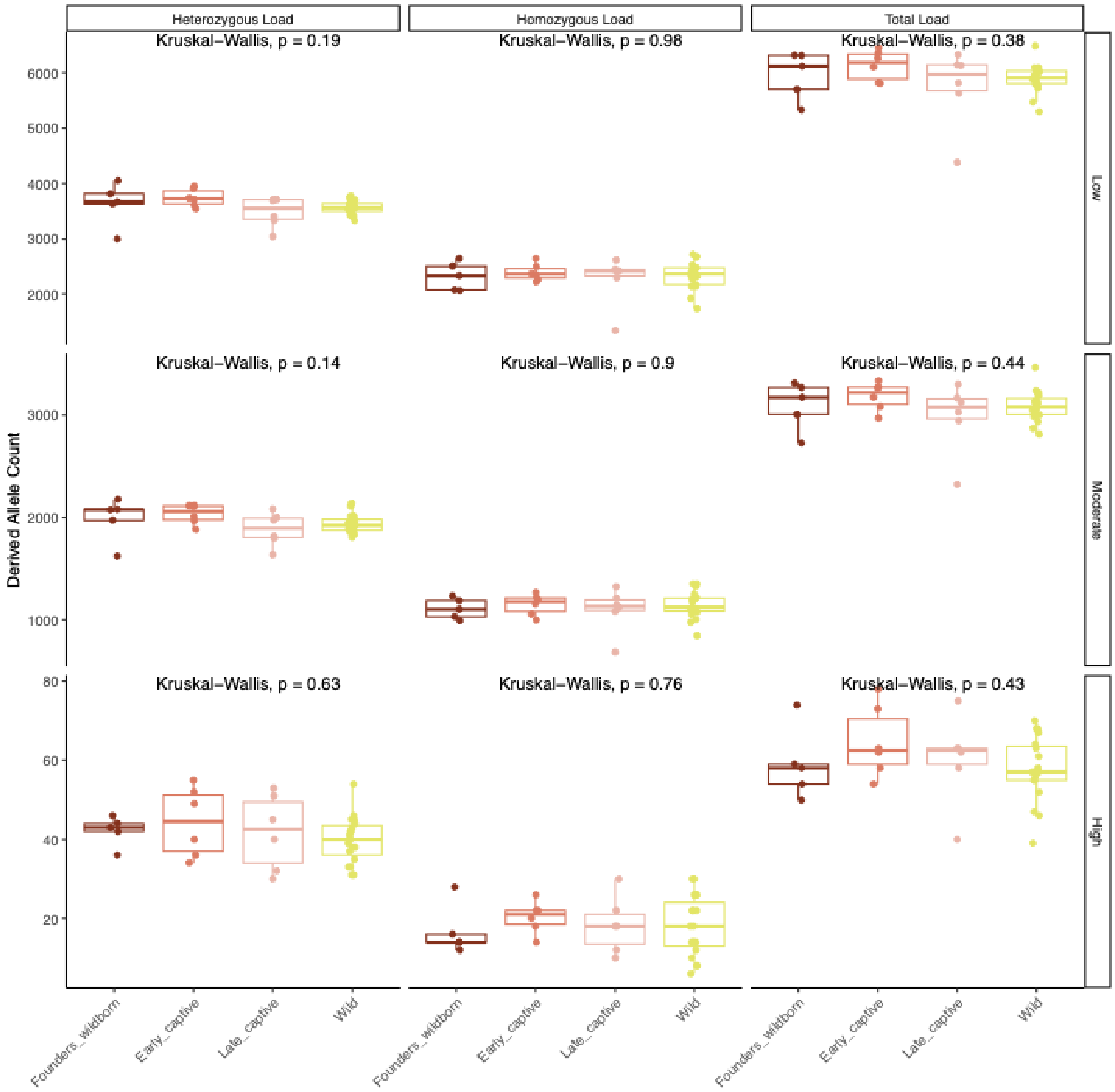

**Figure S12 | Genetic load in modern samples (downsampled to 10x)**. Genetic load is calculated by counting the derived allele in homozygosity, heterozygosity and total load (sum of both) for Low, Moderate and High impact variants. Statistical significance is obtained with a Kruskal-Wallis test for each category.

### Figure S13

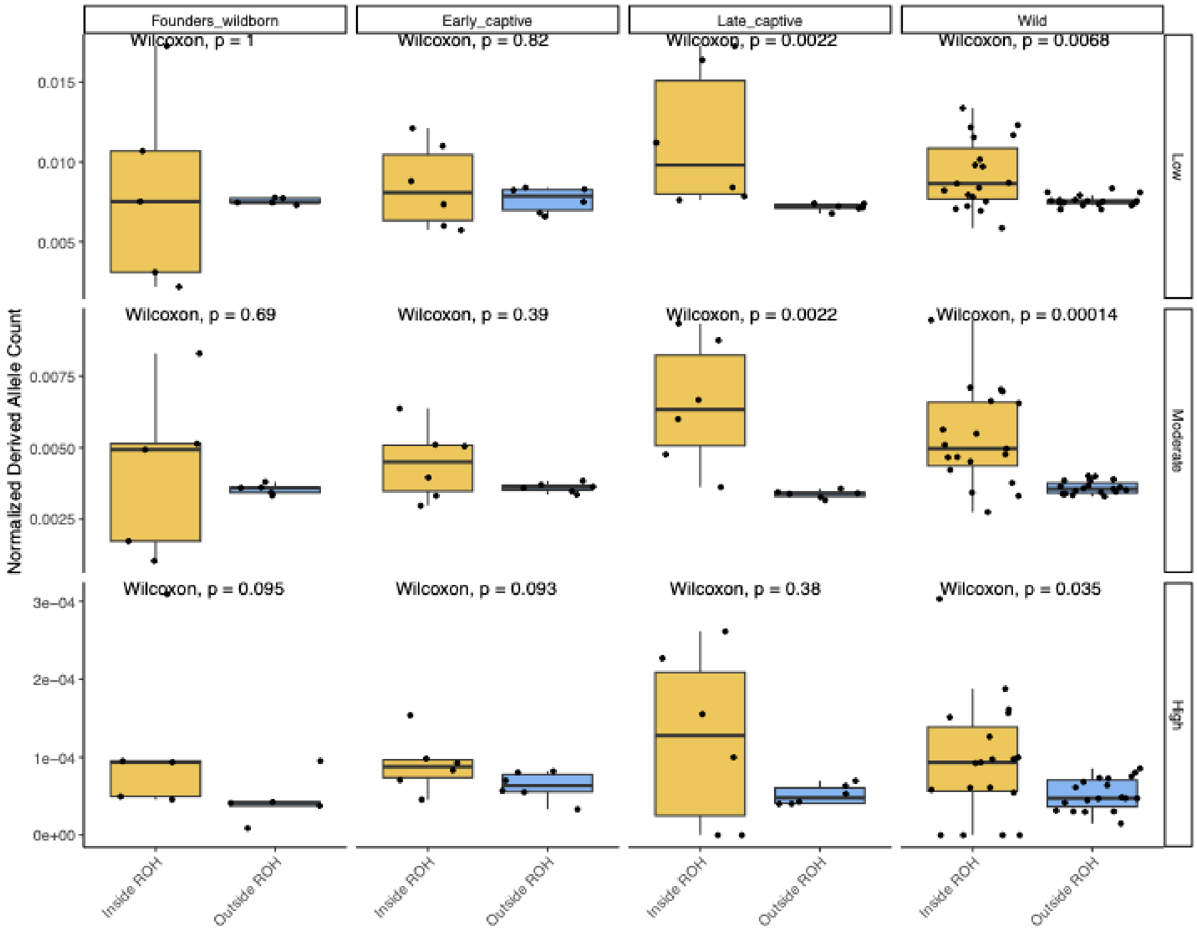

**Figure S13 | Proportion of homozygous load inside and outside ROH** as derived allele counts in homozygosity normalized by the homozygous counts in each category. In all plots, significance statistical tests are calculated with a Wilcoxon rank-sum test.

### Figure S14

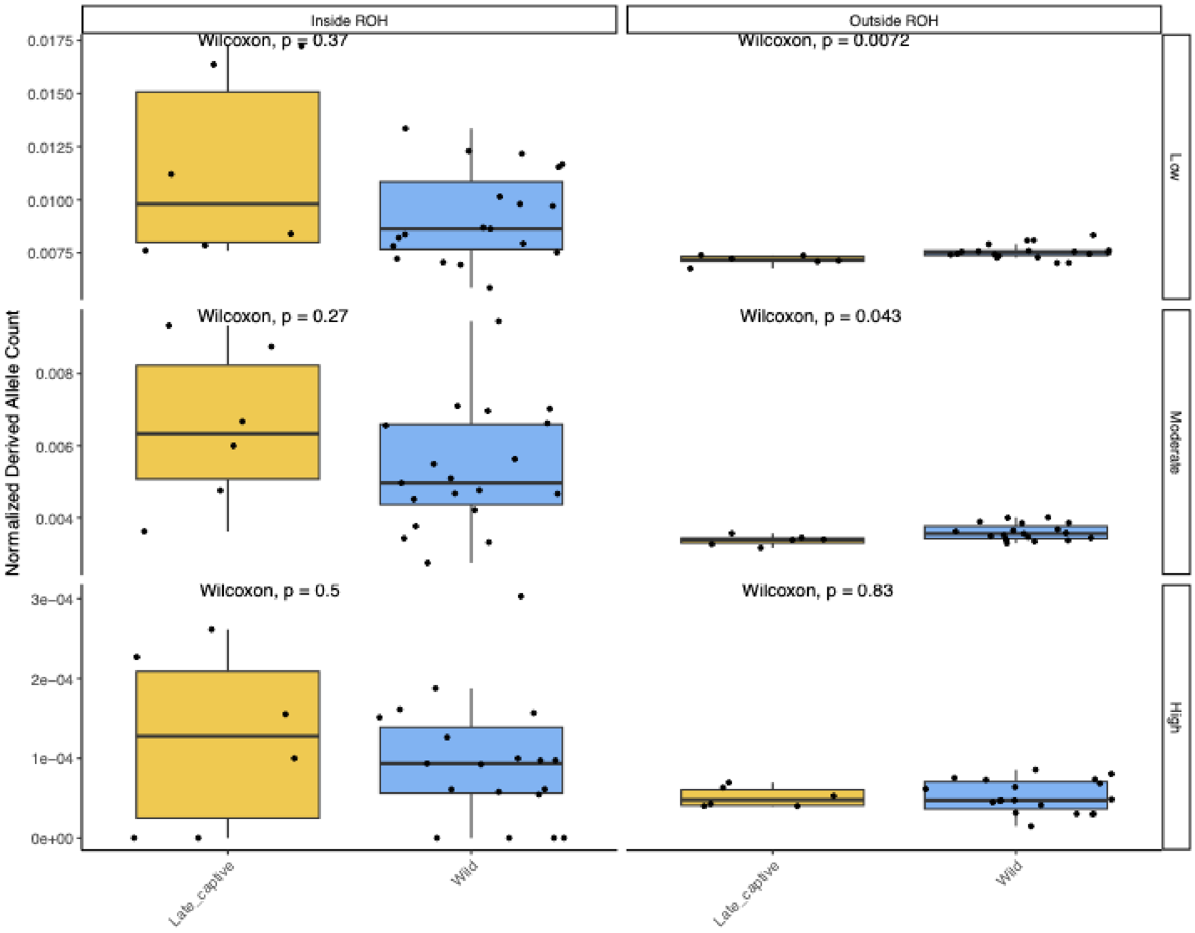

**Figure S14 | Proportion of homozygous load inside and outside ROH** as derived allele counts in homozygosity normalized by the homozygous counts in each category. Only comparing the latest generation in captive breeding (Late_captive) and the wild unmanaged population. In all plots, significance statistical tests are calculated with a Wilcoxon rank-sum test.
